# Genomic surveillance and molecular evolution of fungicide resistance in European populations of wheat powdery mildew

**DOI:** 10.1101/2024.10.30.621051

**Authors:** Nikolaos Minadakis, Jigisha Jigisha, Luca Cornetti, Lukas Kunz, Marion C. Müller, Stefano F.F. Torriani, Fabrizio Menardo

## Abstract

Fungicides are used in agriculture to manage fungal infections and maintain crop yield and quality. In Europe, their application on cereals increased drastically starting from the mid 1970s, contributing to a significant improvement in yields. However, extensive usage has led to the rapid evolution of resistant pathogen populations within just a few years of fungicide deployment. Here we focus on wheat powdery mildew, a disease caused by the ascomycete fungus *Blumeria graminis forma specialis tritici* (Bgt). Previous research on Bgt documented the emergence of resistance to different fungicides and identified various resistance mechanisms. Yet, the frequency, distribution, and evolutionary dynamics of fungicide resistance in Bgt populations remain largely unexplored. In this study we leveraged extensive sampling and whole-genome sequencing of Bgt populations in Europe and the Mediterranean to investigate the population genetics and molecular epidemiology of fungicide resistance towards five major fungicide classes. We analyzed gene sequences and copy number variation of eight known fungicide target genes in 415 Bgt isolates sampled between 1980 and 2023. We observed that mutations conferring resistance to various fungicides increased in frequency over time, and had distinct geographic distributions, likely due to diverse deployment of fungicides across different regions. For demethylation inhibitor fungicides we identified multiple independent events of resistance emergence with distinct mutational profiles, and we tracked their rapid spread in the last decades. Overall, we revealed the evolutionary and epidemiological dynamics of fungicide resistance mutations in European Bgt populations. These results underscore the potential of genomic surveillance and population genetics to enhance our understanding of fungicide resistance.

## Introduction

With more than 130 million tons harvested each year, wheat is among the most important crops in Europe (Eurostat 2024). Starting from the 1970s, the application of fungicides to European wheat fields has become systematic, making Europe the continent with the most intense use of fungicides on cereals (Pimentão et al. 2024). Five main classes of fungicides have been used in the last decades: methyl benzimidazole carbamates (MBCs) and morpholines from the early 1970s, demethylation inhibitors (DMIs) from the late 1970s, quinone outside inhibitors (QoIs) from the 1990s, and succinate dehydrogenase inhibitors (SDHIs) starting in 2002 (Bryson 2022; Jørgensen et al. 2019; Lucas et al. 2015). However, different pathogens have developed resistance to each of these fungicides within a few years from their introduction (Brent 2012; Chartrain and Brown 2023; Jørgensen et al. 2019; Lucas et al. 2015). Because of widespread insensitivity developed by multiple pathogens, the usage of MBCs has declined drastically, while morpholines, QoIs, DMIs and SDHIs continue to be applied on wheat fields. Resistance to these fungicides is managed with mixtures and rotation of products, together with other strategies such as non-chemical control with resistant host varieties (Brent 2012; Corkley et al. 2022; Jørgensen et al. 2019).

Granular up-to-date data about fungicide usage in Europe is not available (Mesnage et al. 2021), however it is known that until 2003, DMIs, morpholines, and QoIs were the three most used fungicides on cereals, while the utilization of SDHIs has increased considerably from the 2010s (Table 5.1.2 in Eurostat 2007; Jørgensen et al. 2019). In addition, there are large differences in fungicide usage across European countries. In Western and Central Europe, fungicide application per hectare of cereal crop has been considerably higher compared to Southern and Eastern Europe (Eurostat 2007). For example, until 2003, fungicide usage on cereals in the UK has been more than 30 times higher than in Italy and four times higher than in Poland (Table 5.1.2 in Eurostat 2007).

In this study we focus on wheat powdery mildew, a disease caused by the biotrophic fungal pathogen *Blumeria graminis forma specialis tritici* (hereafter Bgt). Bgt is considered a pathogen at high risk of developing resistance, because of its short generation time, its ability to recombine sexually, and the very large number of spores produced by any successful infection (McDonald and Linde 2002). Indeed, Bgt populations have quickly become insensitive upon exposure to new fungicides (Jørgensen et al. 2019; Lucas et al. 2015; Vielba-Fernández et al. 2020). For instance, the first Bgt strains insensitive to QoIs were observed just two years after the introduction of this new fungicide class (Chin et al. 2001). Previous research has revealed that in Bgt and other fungi, resistance to QoIs evolved through the G143A mutation in the cytochrome b (*cytb*) gene (Cowger et al. 2022; Fraaije et al. 2002, 2005; Gisi et al. 2002, Sierotzki et al. 2000). Similarly, mutation V295L in the C-14 reductase (*erg24*) gene in the ergosterol pathway has been linked to morpholine resistance (Arnold 2018), while mutations Y136F and S509T in the sterol 14α-demethylase (*cyp51*) gene, along with the presence of multiple copies of this gene, are known to decrease the efficacy of DMIs (Arnold 2018; Arnold et al. 2024; Meyers 2020; Wyand and Brown 2005). Additionally, it was recently proposed that the presence of both alleles for Y136F and S509T in different copies of *cyp51* within the same isolate might be associated with increased resistance to DMIs (Arnold et al. 2024). In contrast, resistance to SDHIs has never been reported in the field (Arnold 2018; FRAC SDHI working group 2024).

Currently, the only systematic monitoring effort to screen the sensitivity of Bgt populations to different fungicides in Europe is undertaken by the private sector. Every year the Fungicide Resistance Action Committe (FRAC) gathers data generated by fungicide producers. However, only a summary of the results and usage recommendations are made available to the public. From this information we know that Bgt populations are at least partially insensitive to QoIs, DMIs and morpholines in Northern Europe, and that in Western Europe Bgt is particularly resistant to DMIs (FRAC QoI Working Group 2024; FRAC SBI Working Group 2024).

Here we take advantage of a large, previously published collection of genome sequences from Bgt isolates sampled in Europe and surrounding regions (Jigisha et al. 2024) to perform a systematic screening for fungicide resistance mutations. We focus on eight known Bgt genes coding for proteins targeted by the five fungicide classes described above. We investigate variation in the amino acid sequence and copy number of these target genes, with particular attention to mutations that are known to confer resistance. We determine their prevalence in different regions, and we track how they spread through time and space. We found that resistant mutations for different fungicides have distinct geographic ranges, likely a consequence of different fungicide usage in different countries. Moreover, we identify novel amino acid mutations in *erg24* and we suggest that they are likely to confer resistance to morpholines. Finally, we show that mutations conferring resistance to DMIs are abundant in Europe and Turkey, and that resistance to DMIs evolved multiple times independently and spread rapidly throughout the continent.

## Results

To monitor the prevalence of fungicide resistance mutations, and to study how Bgt populations evolved in response to fungicide pressure, we used whole-genome sequencing data of 415 *Bgt* isolates obtained from Jigisha et al. (2024). That study reported that wheat powdery mildew epidemics in Europe and the Mediterranean can be divided in five populations occupying different geographic ranges: one population in Northern Europe (population N_EUR), two populations in Southern Europe (populations S_EUR1 and S_EUR2), one population in Turkey (population TUR), and one in the Middle East (population ME; see Figure 1 in Jigisha et al. 2024).

**Figure 1.**
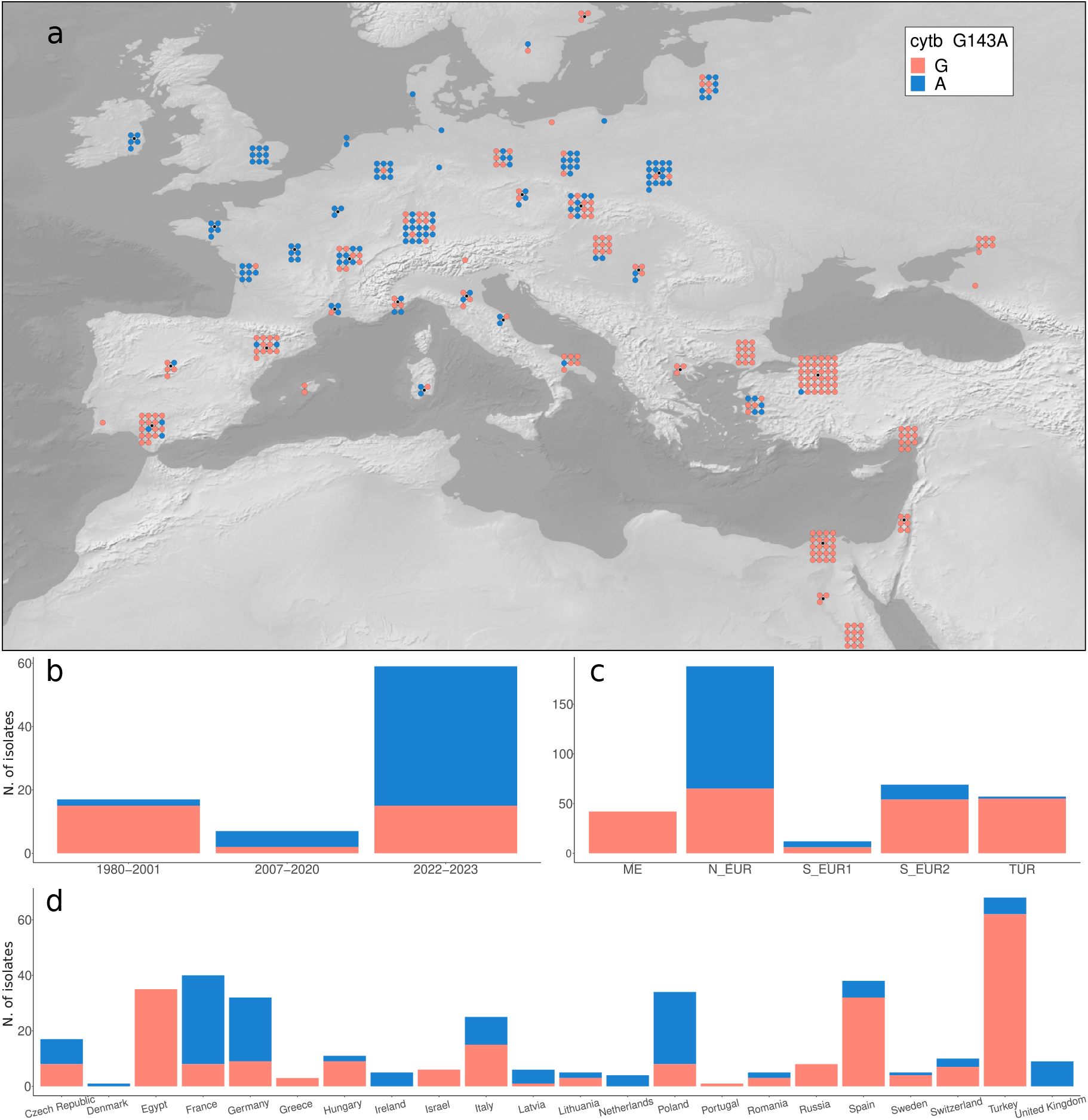
*cytb* mutation G143A. **(a)** Geographic distribution of G143A. **(b)** Frequency of G143A by year of collection (*temporal* dataset). **(c)** Frequency of G143A by population. **(d)** Frequency of G143A by country of origin.

Here, we defined three datasets: 1) a set of 415 *Bgt* isolates collected in Europe and the Mediterranean between 1980 and 2023 (dataset *Europe+*), 2) a subset of *Europe+* with 368 isolates collected between 2015 and 2023 (dataset *Europe+_recent*) and 3) a subset of 83 samples collected over a period of more than 40 years from the UK, France and Switzerland (*temporal* dataset; Supplementary Data S1). We selected these countries for the *temporal* dataset because we wanted to study shifts in fungicide resistance over multiple decades, and these were the only countries with consistent sampling over the years. For most of the analyses, and unless otherwise stated, we used the *Europe+_recent* dataset. We generated amino acid sequence alignments for the eight known fungicide targets that were the focus of this study (Table 1; Experimental procedures). For the three subunits of the succinate dehydrogenase gene (*sdhB, sdhC*, and *sdhD*; targets of SDHI fungicides), the C-8 sterol isomerase (*erg2*; target of morpholines), and beta tubulin (*Btub*; target of MBC fungicides), we only found a few amino acid mutations at low frequency (between 0.3% and 2.5%; Table S1). Moreover, there was virtually no variation in the copy number of these genes (Figure S1). Thus, we focused on *cytb, erg24* and *cyp51* for the rest of the study.

**Table 1.**
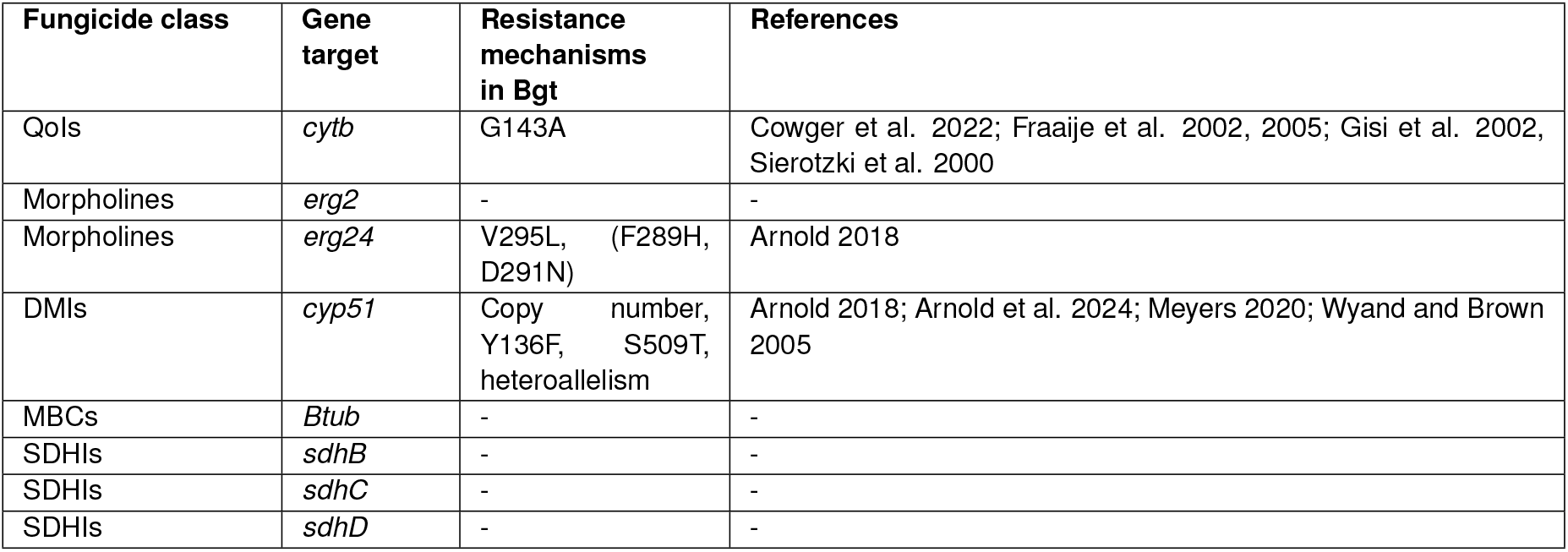
Investigated gene targets. The eight gene targets investigated in this study and the corresponding fungicide classes. Observed resistance mechanisms are shown, novel potential resistance mutations that were never observed in Bgt are reported in parenthesis.

| Fungicide class | Gene target | Resistance mechanisms in Bgt | References |
| --- | --- | --- | --- |
| QoIs | *cytb* | G143A | Cowger et al. 2022; Fraaije et al. 2002, 2005; Gisi et al. 2002, Sierotzki et al. 2000 |
| Morpholines | *erg2* | - | - |
| Morpholines | *erg24* | V295L, (F289H, D291N) | Arnold 2018 |
| DMIs | *cyp51* | Copy number, Y136F, S509T, heteroallelism | Arnold 2018; Arnold et al. 2024; Meyers 2020; Wyand and Brown 2005 |
| MBCs | *Btub* | - | - |
| SDHIs | *sdhB* | - | - |
| SDHIs | *sdhC* | - | - |
| SDHIs | *sdhD* | - | - |

### *cytb* – QoI resistance

Systematic data about QoI resistance in European Bgt populations is lacking, however wheat powdery mildew in Europe is considered largely insensitive to these fungicides (Cowger et al. 2022; FRAC QoI working group 2024; Jørgensen et al. 2019; Lucas et al. 2015). To investigate the sensitivity of European populations of Bgt to QoI fungicides we focused on *cytb*, a mitochondrial gene coding for a subunit of cytochrome bc_1_, which is known to be the single target of QoIs. The amino acid substitution G143A is associated with field resistance to QoI fungicides in many fungal species, including Bgt (Cowger et al. 2022; Dorigan et al. 2023). Across all isolates we found one single nucleotide mutation (G428C), corresponding to the amino acid change G143A. This mutation was present in 146 out of 368 (39.7%) isolates of the *Europe+_recent* dataset (Figure 1a, Table S1), suggesting that resistance to QoIs is widespread but not complete in European Bgt populations.

However, there were large differences in the prevalence of G143A in different geographic regions. Notably, in Northwestern Europe almost all isolates carried the resistant allele. In Central and Eastern Europe, G143A was present at intermediate frequencies, while in the south of Europe it was rare, and in the Middle East it was completely absent (Figure 1). We also investigated how resistance to QoIs changed over time in Western Europe by comparing samples collected in three different periods: 1980-2001, 2007-2020, and 2022-2023 (*temporal* dataset). We observed an upward trend in the occurrence of G143A (Fisher’s exact test *p*-value < 0.001 for the comparison between 1990-2001 and 2022-2023 in the *temporal* dataset, Figure 1b; Table S1), indicating an expansion of QoI resistance in the last two decades in Western Europe. Altogether, these analyses showed that resistance to QoIs is almost complete in Western Europe, a region in which QoIs have been used intensely in the past, and that G143A is also abundant in Eastern Europe, reflecting the broader application of this class of fungicides.

### *erg24* – morpholine resistance

Morpholine fungicides have been used on cereals since the 1970s, although they have been less popular than DMIs and their deployment has decreased over time (Eurostat 2007; Jørgensen et al. 2019). Reduced Bgt sensitivity to morpholines was reported in different European countries (Arnold 2018; Godet and Limpert 1998; Jørgensen et al. 2019; FRAC SBI working group 2024; Zziwa 1999), but as for other fungicides there is no public data about the prevalence of morpholine resistance. Morpholines are inhibitors of ergosterol biosynthesis, and their main targets are *erg2* -which showed little amino acid variation in our dataset- and *erg24* (Chartrain and Brown 2023). A previous study on a panel of Bgt samples collected in England reported two widespread mutations in *erg24*: V295L, which was present exclusively in resistant isolates and was shown to confer resistance to fenpropimorph (a morpholine-derived fungicide) in yeast, and Y165F which was not associated with fungicide resistance (Arnold 2018). In addition, the same study reported that the mutation D291N was associated with morpholine resistance in barley powdery mildew, but it was not observed in Bgt.

In our dataset (*Europe+*), all but one isolate carried a single copy of *erg24* (Figure S1). We identified a total of eight amino acid mutations, three of which were previously observed: V295L, Y165F, and D291N (which we report for the first time in Bgt). The five new ones included D137E, F289H, and three additional mutations found in a single strain, which were therefore excluded from further analyses (Table 2, Table S1).

**Table 2.** Summary of *erg24* amino acid mutations. Only amino acid mutations that are present in more than one isolate are shown. For the geographic distribution of each mutation see Figure 2, and Figures S2-S6. The results of the test for increased frequency over time are reported in Table S1. (*: *p*-value < 0.1). For the previous evidence about resistance see Arnold 2018. Bh: *Blumeria hordei* (barley powdery mildew).

| Mutation | N. of isolates | Geographic distribution | Increase in frequency over time | Previous evidence about resistance |
| --- | --- | --- | --- | --- |
| V295L | 141 | Northern Europe | Yes* | Yes |
| Y165F | 74 | widespread | No | No |
| F289H | 36 | Northern Europe | Yes | No |
| D137E | 17 | Southern Europe | - | No |
| D291N | 8 | Northern Europe | - | Yes in Bh |

V295L was the most frequent mutation in the *Europe+_recent* dataset (141/368 - 38%) and was present almost exclusively in isolates from Northern Europe, with an especially high prevalence in the west (Figure S2). We also found an increase in the proportion of isolates having this mutation in Western Europe over time (Fisher’s exact test *p*-value = 0.075 for the comparison between 1990-2001 and 2022-2023 in the *temporal* dataset; Figure S2, Table S1).

V295L is located within a sterol binding pocket where both the substrates of *erg24* and the morpholine fenpropimorph are predicted to dock. It was hypothesized that fenpropimorph can compete with the sterol substrate and block it from binding *erg24*, and that mutations in or nearby this domain might decrease the affinity between *erg24* and fenpropimorph (Arnold 2018). We found two mutations in close proximity of V295L: 1) D291N was present in 8 isolates sampled in 2022 and 2023 in Northern and Central Europe, and it is associated with morpholine resistance in barley powdery mildew (Figure S3), 2) F289H was present in 36 isolates from the Northern European population (N_EUR) and was not present in samples collected before 2007 (Fisher’s exact test *p*-value = 0.195 for the comparison between 1990-2001 and 2022-2023 in the *temporal* dataset; Figure S4, Table S1). In contrast, the frequency of the Y165F mutation did not change over time, and we did not find any pattern in its geographic distribution (Fisher’s exact test *p*-value = 0.732 for the comparison between 1990-2001 and 2022-2023; Figure S5, Table S1). Finally, the newly discovered mutation D137E was present in 17 Southern European isolates (Figure S6).

To investigate the evolution of *erg24* in Europe, we inferred a haplotype network based on nucleotide sequences (Figure 2). At the protein level, the wildtype (no amino acid substitutions) was the most common haplotype, and was especially abundant in Southern Europe, Turkey, and the Middle East. The second most frequent haplotype contained only the V295L mutation while the third was the Y165F + V295L haplotype, both almost exclusively present in Northern Europe. The haplotype containing only Y165F was probably derived independently from a wildtype haplotype and only present in Southern Europe and Turkey. Finally, the mutation F289H was present only in the absence of other mutations, except for one isolate with the V295L + F289H haplotype (Figure 2c).

**Figure 2.**
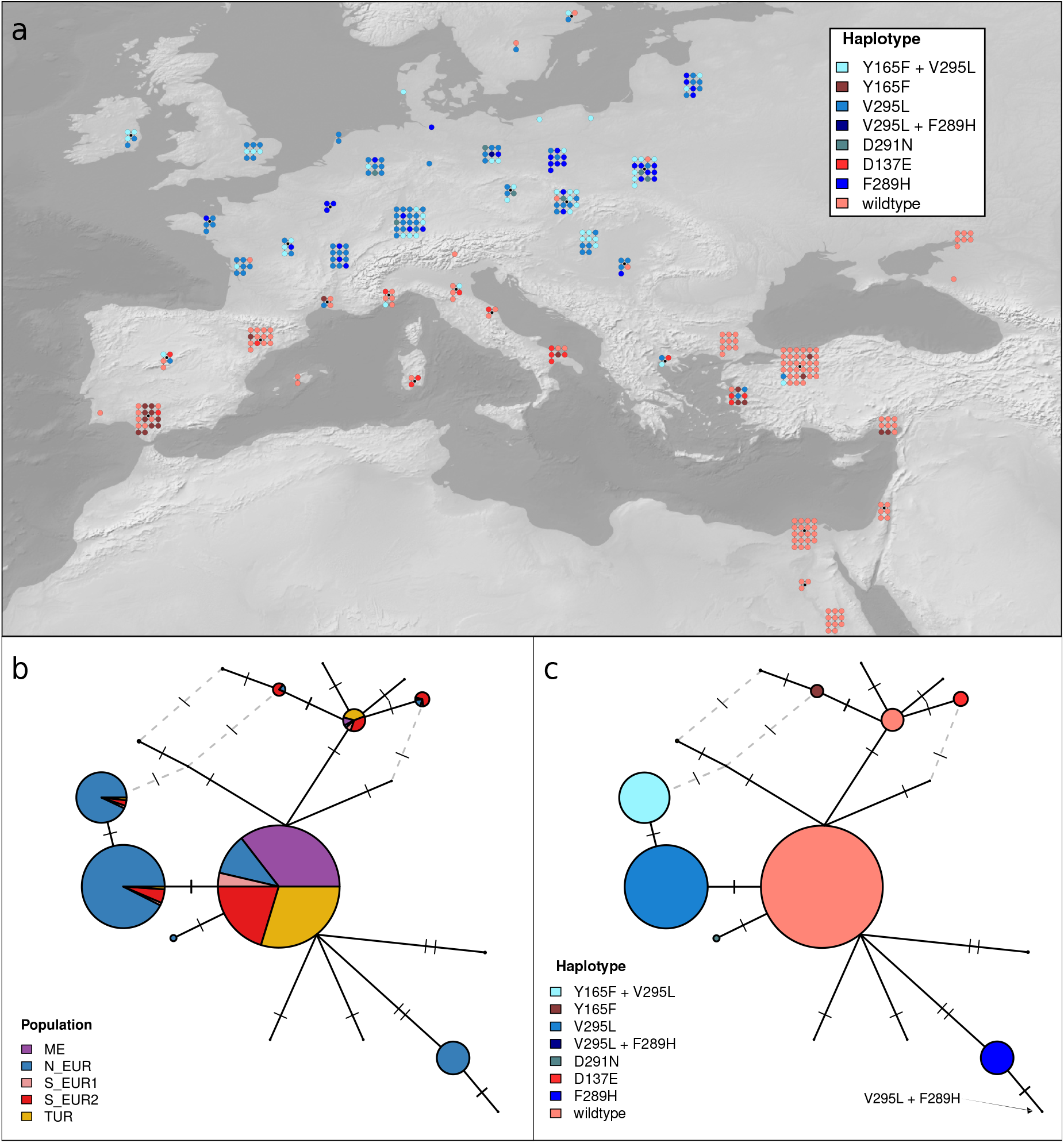
Haplotype network of *erg24*. Networks were inferred based on nucleotide sequences, while haplotypes are defined based on the five most common amino acid mutations. Each node of the network represents a unique nucleotide sequence. The size of nodes is proportional to the number of isolates in which that nucleotide sequence was observed. Ticks on edges connecting different nodes represent the number of nucleotide differences. Dashed grey edges represent alternative connections. Alternative connections with more than two nucleotide differences are not plotted. **(a)** Geographic distribution of *erg24* haplotypes. Putatively resistant haplotypes are represented as blue shades, putatively susceptible haplotypes are depicted with red shades **(b)** Haplotype network of *erg24* colored by population. **(c)** Haplotype network of *erg24* colored by haplotype (colors as in panel **a**).

### *cyp51* - DMI resistance

DMIs are another important component of disease control on wheat in Europe, with most fields receiving between one and three applications per season (Jørgensen et al. 2019). Resistance to DMIs was observed in different pathogens and is widespread on the continent (FRAC SBI working group 2024). In Bgt, there are three known mechanisms providing resistance to DMIs all occurring in the *cyp51* gene: copy number variation and the amino acid mutations Y136F and S509T (Arnold 2018; Arnold et al. 2024; Meyers 2020; Wyand and Brown 2005). In addition, a recent study proposed that “heteroallelism” for Y136F and S509T could also be associated with increased insensitivity to DMIs (Arnold et al. 2024).

### Copy number variation

An increased number of copies of *cyp51* leads to reduced sensitivity to DMIs by increasing the expression level of the target gene (Arnold et al. 2024; Meyers 2020). We estimated the number of copies of *cyp51* for each isolate by comparing the gene average coverage with the genome-wide average coverage (see Experimental procedures). We found between one and seven copies, with 88% (324/368) of the isolates carrying more than one copy of *cyp51* (Supplementary Data S1). The individuals carrying one copy were mostly sampled in Southern Europe and the Middle East (ME and S_EUR2 populations; Figure 3). We also observed that there was no increase in the average number of copies between isolates collected before 2001 and isolates collected in 2022-2023 (Welch two sample t-test *p*-value = 0.351; Figure 3b). To confirm our estimates of the number of *cyp51* copies, we used the previously published PacBio genome assembly of one of the isolates in our collection (CHVD042201; Kunz et al. 2024), which we estimated to have four copies. We identified four identical copies of *cyp51* in the PacBio assembly of this isolate, organized in a tandem repeat pattern (Figure S7). Furthermore, as independent proof that our copy number estimation was accurate, we identified two isolates (94202 and JIW11) that were included in another study in which their copy number was estimated with a ddPCR approach (Arnold et al. 2018). Both the coverage ratio and the ddPCR results suggested that the two isolates have two copies of *cyp51* each, further supporting the robustness of our results.

**Figure 3.**
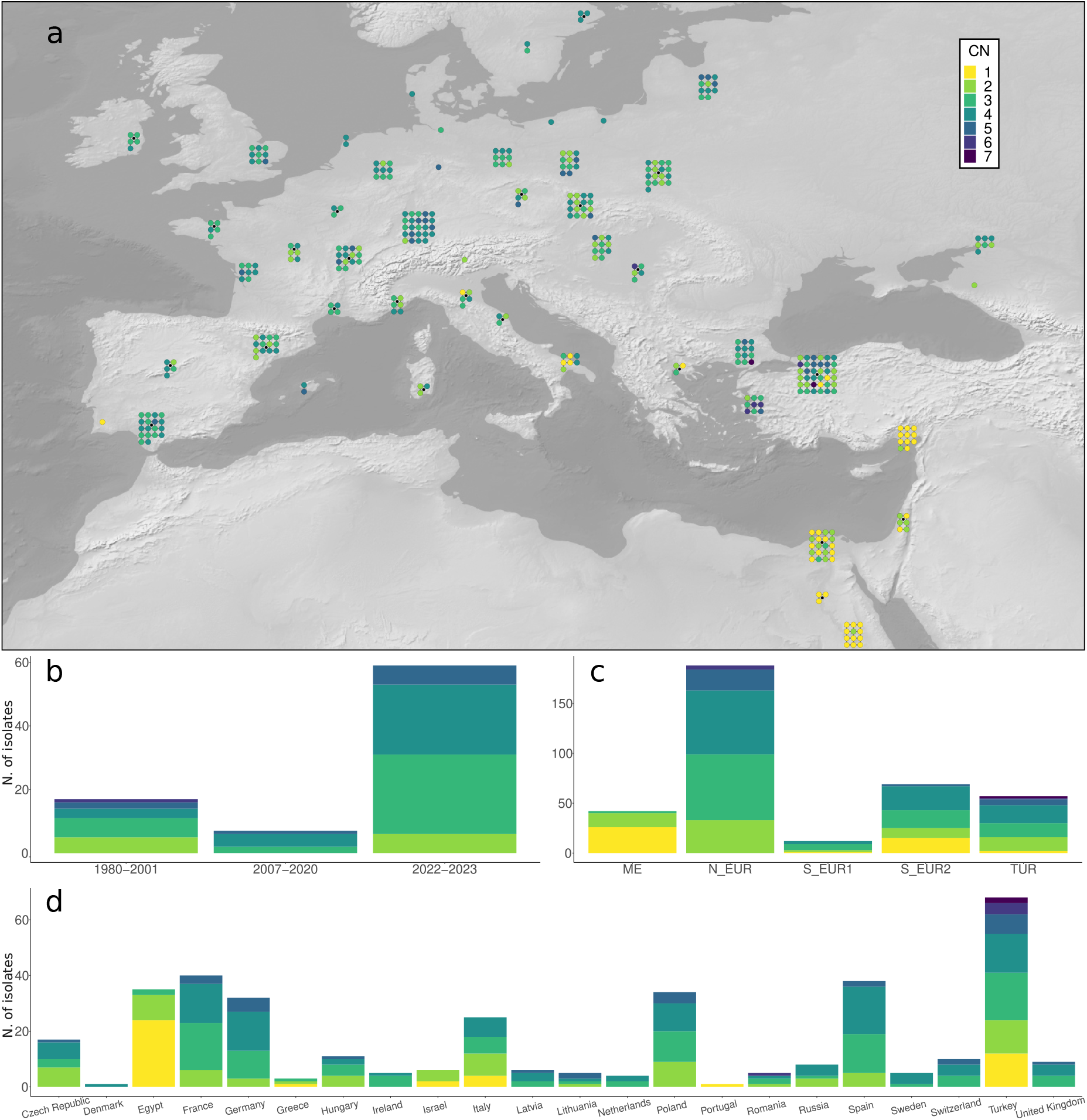
*cyp51* copy number variation. **(a)** Geographic distribution of the number of copies (CN) of *cyp51* per isolate. **(b)** Number of *cyp51* copies by year of collection (*temporal* dataset). **(c)** Number of *cyp51* copies by population. **(d)** Number of *cyp51* copies by country of origin.

### Amino acid mutations in cyp51

Due to the variable copy number of *cyp51* we could not resolve the sequence of each copy with our short read data. However, we could identify mutations present in at least one copy of *cyp51* for each sample, though not necessarily in all copies (see Experimental procedures). With this approach we detected eight amino acid mutations, one of which was present in only one isolate and was not considered further (Table S1).

The most abundant *cyp51* mutation was Y136F (298/368, 81%; Table S1), which is known to confer partial resistance to DMI fungicides in many crop pathogens including Bgt (Arnold 2018; Arnold et al. 2024; Dorigan et al. 2023; Meyers 2020; Wyand and Brown 2005). Almost all isolates in Northern Europe, Spain and Turkey carried this mutation, while it was less abundant in Central and Southern Italy, and in the Middle East. Notably, Y136F was only found in isolates with more than one copy of *cyp51*, and its frequency increased with higher copy number (Figure S8). Moreover, we observed a significant increase in frequency of Y136F over time (Fisher’s exact test *p*-value < 0.001 for the comparison between 1980-2001 and 2022-2023 in the *temporal* dataset; Figure S8, Table S1).

The other amino acid mutation that is known to be associated with resistance to DMIs is S509T, which was previously described for wheat (Arnold 2018; Arnold et al. 2024) and barley powdery mildew (Tucker et al. 2020; Zulak et al. 2018). S509T confers resistance to some DMIs and might also act as a compensatory mutation to restore the lower fitness conferred by the Y136F mutation in absence of fungicides (Arnold 2018; Arnold et al. 2024; Cools et al. 2011). We found S509T in 62 of 386 samples (16%), mostly in Northern Europe, and we observed that its earliest presence in the whole dataset was in Turkey in a sample collected in 2018, pointing to a recent increase in frequency (Fisher’s exact test *p*-value = 0.031 for the comparison between 1990-2001 and 2022-2023 in the *temporal* dataset; Figure S9, Table S1). Notably, S509T co-occurred with Y136F in more than 90% of the isolates carrying the mutation (57/62).

We observed two additional mutations at high frequency, S79T (57%) and K175N (64%; Figures S10-S11, Table S1). These mutations were previously detected in Bgt, but they could not be associated with resistance (Wyand and Brown 2005). S79T and K175N were especially abundant in Spain and Turkey, but also in many countries in Northern Europe. They never co-occurred with S509T, and they were never detected together except in the presence of Y136F. Indeed, the combination Y136F + S79T + K175N was the most frequent in the *Europe+_recent* dataset (57%). The three remaining mutations are not known to be associated with fungicide resistance, and they were found at low frequency. T271S and L236F occurred together in the same isolates in Egypt and Greece (Figures S12-S13), while I3K was previously described in the UK, but was mostly found in the Middle East (Figure S14).

### Recent evolution of cyp51 in Europe

Since it was impossible to resolve the multiple copies of *cyp51* we could not study this gene at the haplotype level like we did for *erg24*. Therefore, we used an alternative method to investigate the recent evolution of this locus. We inferred identical-by-descent (IBD) segments between pairs of Bgt isolates using IsoRelate (Henden et al. 2018). IBD segments are large chromosomal sections which are nearly identical between two individuals because they have been inherited from a recent common ancestor (roughly within 25 sexual generations; see Experimental procedures). In a previous study Bgt populations in Europe and Turkey showed an excess of IBD pairs over *cyp51*, indicating recent positive selection for this locus (Jigisha et al. 2024). Here we identified all pairs of isolates which are IBD over the *cyp51* locus and grouped them into clusters. Large clusters with many IBD isolates over *cyp51* represent instances in which the locus has been inherited by many individuals over several generations, indicating a potential fitness advantage.

We identified two large clusters (clusters **i** and **ii** in Figure 4; Supplementary Data S1), which comprised 129 and 75 isolates from Europe and Turkey. The rest of the samples were grouped in smaller clusters or were not IBD with any other strain over the *cyp51* locus. The two main clusters had different mutational profiles (Figure 4). Furthermore, isolates in cluster **i** had more copies of *cyp51* compared to cluster **ii** (Welch two sample t-test *p*-value < 0.001; Figure 4b; Supplementary Data S1). This analysis suggests that resistance to DMIs emerged at least twice independently and was achieved through higher copy number and the Y136F mutation. Among the two largest clusters, S509T emerged only in cluster **ii**, while S79T and K175N emerged only in cluster **i**, though we do not know whether the latter two mutations affect the fitness of isolates carrying them.

**Figure 4.**
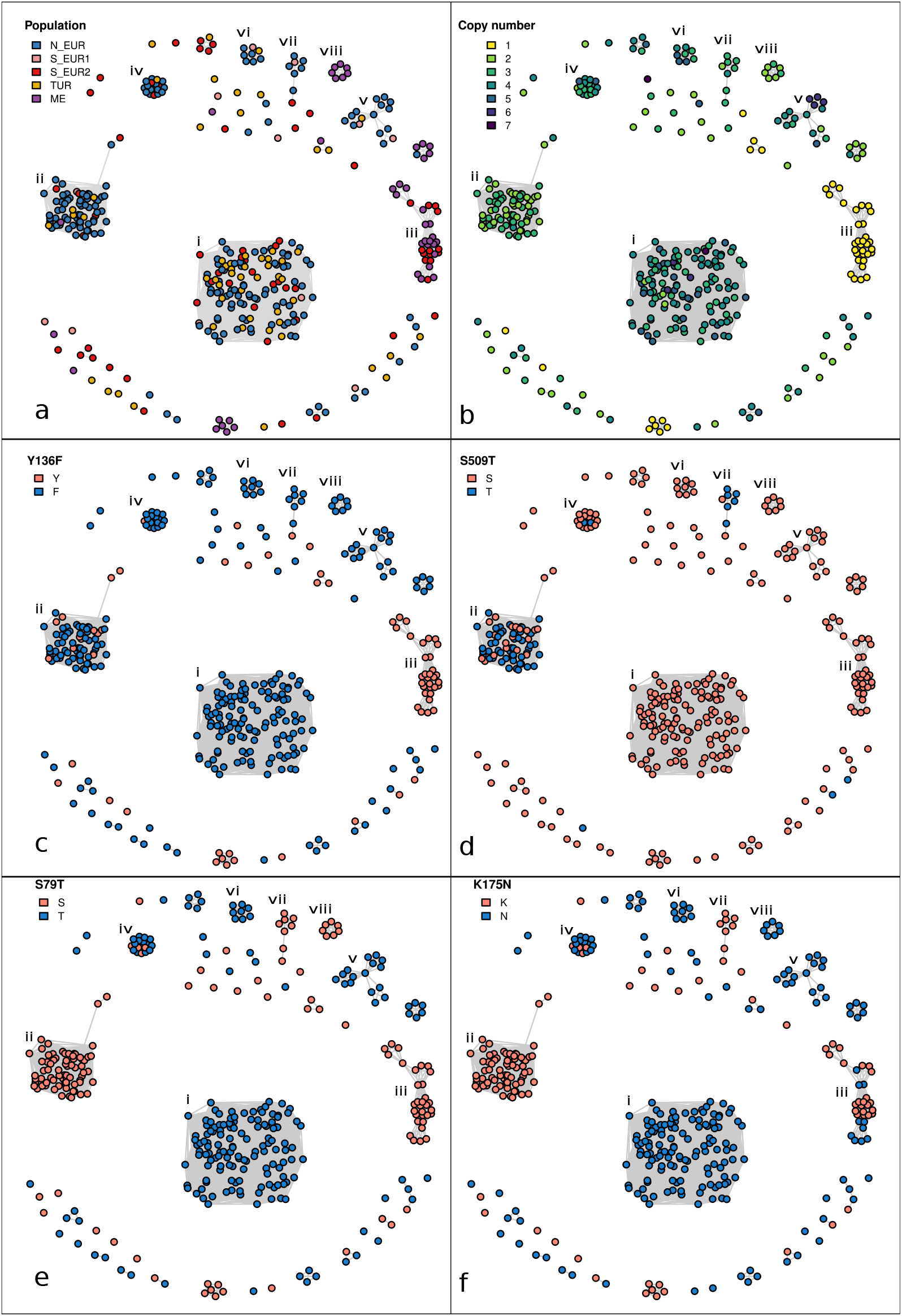
Identical-by-descent (IBD) clusters for *cyp51* locus. Relatedness network for the *cyp51* locus. Each node represents one isolate (368 samples belonging to the *Europe+_recent* dataset), edges connect isolates that are identical-by-descent over the *cyp51* locus. Isolates are colored by: **(a)** population **(b)** copy number **(c)** amino-acid mutation Y136F **(d)** amino-acid mutation S509T **(e)** amino-acid mutation S79T and **(f)** amino-acid mutation K175N. The eight largest clusters are labeled using latin numerals.

Furthermore, by inspecting the heterozygosity of the mutations we found that all *cyp51* copies in all isolates belonging to cluster **i** were identical (see Experimental procedures). This indicates that the *cyp51* mutations likely preceded the first gene duplication, or alternatively that *cyp51* was duplicated before the mutations occurred, with recurring gene conversion events making all copies identical in all isolates. Conversely, in cluster **ii**, at position 136 (Y136F mutation), 72 out of the 75 isolates had different “alleles” in different copies, while the remaining three isolates had either Y (two isolates), or F (one isolate) at all copies. Similarly, in position 509 (S509T mutation), 65 isolates were “heteroallelic”, while two had T and eight S at all copies. Thus, it is likely that in cluster **ii**, Y136F and S509T emerged after the first duplication.

To further investigate the two main clusters, and to confirm that they represent independent emergence events, we analyzed the structure of the *cyp51* locus. We have already shown that *cyp51* is present in four identical copies due to repeated tandem duplication events in CHVD042201 (the isolate with an available long reads assembly). CHVD042201 belongs to cluster **i**, and we used the normalized coverage over the *cyp51* locus to estimate the length of the unit of the tandem repeat in all isolates (Figure 5; see Experimental procedures). We found that all isolates belonging to cluster **i** had the same tandem repeat unit, about 2.5 Kb in length. For cluster **ii** we could not identify the exact length of the unit, as it extended in a genomic region with repetitive sequences. Nonetheless, we found that for all cluster **ii** members the tandem repeat unit was longer than for cluster **i**, and had different start and end points (Figure 5). Overall, this analysis confirmed that isolates in the same cluster have related *cyp51* loci and that the two clusters likely represent independent emergence events.

**Figure 5.**
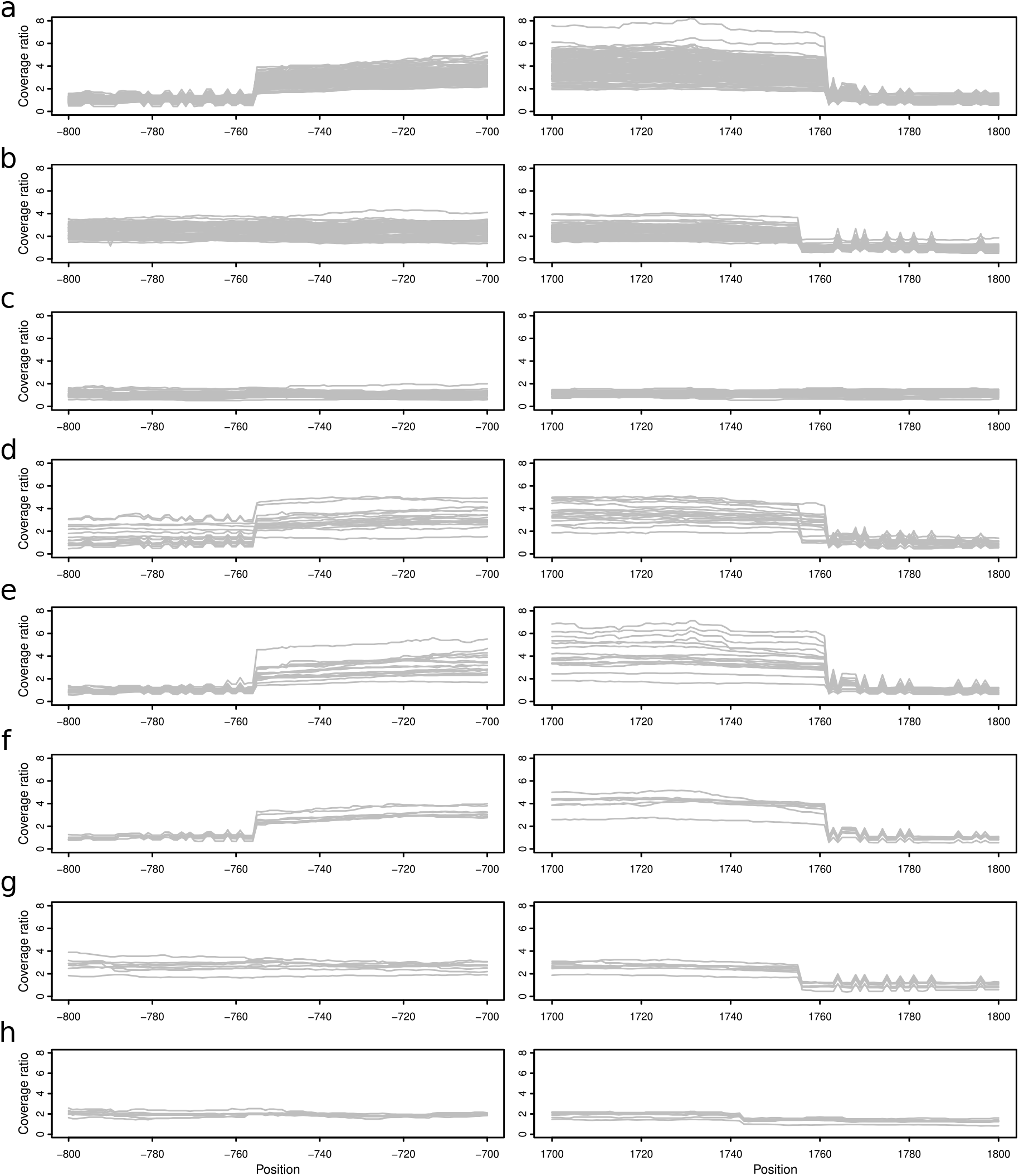
Tandem repeat unit at *cyp51* locus. Normalized coverage around the *cyp51* locus for each isolates belonging to the eight largest clusters as defined in Figure 4 **(a)** cluster **i (b)** cluster **ii (c)** cluster **iii (d)** cluster **iv (e)** cluster **v (f)** cluster **vi (g)** cluster **vii (h)** cluster **viii**). Each gray line represents one isolate. The x axis represents the genomic coordinates in base pairs relative to the beginning of *cyp51* (because *cyp51* is in an inverted orientation this corresponds to the last nucleotide of the stop codon).

The geographic distribution of the samples indicates that upon the evolution of different resistant genotypes both clusters expanded rapidly. This was true especially for cluster **i**, as it was well represented in the two European populations (N_EUR and S_EUR2) and in Turkey (TUR; Figure 4a). Both clusters are now widespread wherever DMIs are used in Europe and in Turkey (Eurostat 2007; Jørgensen et al. 2019; Lucas et al. 2015; Ölmez et al. 2023), but not in the Middle East, where DMIs have not been adopted.

## Discussion

Monitoring fungicide sensitivity in pathogen populations is a crucial aspect of resistance management. Once the molecular mechanism of fungicide resistance is known, monitoring can be undertaken with molecular methods (Yin et al. 2023). With this approach, and similarly to what has previously been done for other pathogens or other geographic regions (Arnold et al. 2024; Cook et al. 2021; Cowger et al. 2022; Estep et al. 2015; Ölmez et al. 2023; Torriani et al. 2009), we characterized fungicide resistance in wheat powdery mildew populations in Europe.

***cytb* - QoI resistance**

The mutation G143A in *cytb* is the only known path to QoI resistance in Bgt, and genetic screening is commonly used instead of sensitivity testing (Arnold 2018; Cowger et al. 2022; FRAC QOI working group 2024; Meyers 2020). Our results revealed a gradient in the frequency of G143A in Northern Europe. All samples from Ireland, England, Northern France, and the Netherlands carried the resistant allele, while its frequency decreased towards the east and the south (Figure 1). This pattern is comparable to what was reported for the same mutation in *Zymoseptoria tritici* (Lucas et al. 2015). The intermediate frequency of G143A in Bgt samples from Central and Eastern Europe could be due to two factors: 1) gene flow from west to east following the main wind direction, similar to what was reported for *Zymoseptoria tritici* (Torriani et al. 2009). This mechanism could result in intermediate levels of G143A in Eastern Europe independently of selective pressure acting on the mutation (i.e., independently of QoIs applications). 2) A late adoption of QoIs in Central and Eastern Europe, with a less intense utilization compared to Northwestern Europe, and therefore a lower fitness advantage of G143A. Finally, we found G143A at low frequency in Southern Europe and Turkey, possibly due to a mix of migration from Northern Europe and low-level selection. However, without up-to-date data about QoI applications it is hard to disentangle the effects of migration and selection on the prevalence of G143A in different regions.

### *erg24* - Morpholine resistance

While morpholines are less important than QoIs and DMIs, they are still used broadly, and their performance in the field is considered good (FRAC SBI working group 2024; Jørgensen et al. 2019; Lucas et al. 2015). Moreover, until at least 2003 they were the third most used fungicide on cereals in Europe, although with a decreasing trend (Eurostat 2007). Our data shows that V295L in *erg24*, the only mutation known to be associated with resistance to morpholines in Bgt, is ubiquitous in Northern Europe, and absent elsewhere. Similarly to G143A in *cytb*, we observed a gradient in the frequency of V295L in Northern Europe, with 100% frequency of V295L in Ireland and the UK, decreasing toward the south and the east (Figure S2). This likely reflects the more intense utilization of morpholines (and fungicides in general) in Northwestern Europe, although when interpreting the frequency of V295L we must apply the same caveat discussed for G143A in *cytb*.

Beyond V295L, we suggest that F289H and D291N are likely to confer resistance to morpholines. This conclusion is based on multiple pieces of evidence: 1) both mutations are located in close proximity of V295L, which was shown to be associated with morpholine resistance (Arnold 2018). 2) Both mutations were never sampled before 2007, and they are found exclusively in Northern Europe, suggesting that their frequency might have increased because of recent selective pressure caused by morpholines. 3) D291N was found to be associated with resistance in barley powdery mildew (Arnold et al. 2018). To test this hypothesis, we encourage further investigations of these mutations, for example by combining sensitivity testing and sequencing of *erg24* of large contemporary Bgt populations. If F289H and D291N were to be confirmed to be resistance-conferring mutations, it would mean that morpholine resistance emerged multiple times through different mutational paths.

### *cyp51* - DMI resistance

In Europe, DMIs are the most common fungicides to control wheat pathogens. Insensitivity to DMIs is widespread, and in Bgt is achieved with three known mechanisms: multiple copies of *cyp51*, which increase the expression level of the gene, and the mutations Y136F and S509T (Arnold 2018; Arnold et al. 2024; Meyers 2020; Wyand and Brown 2005). Moreover, a fourth potential mechanism was proposed recently: “heteroallelism” at positions 136 and 509 (Arnold et al. 2024). As for other resistance-conferring mutations, S509T was mostly present in Northern Europe (Figure S9). Conversely, the distribution of strains with more than one copy of *cyp51* and the distribution of Y136F were broader and very similar (Figure 3 and Figure S8). Almost all strains in Northern Europe, Spain, and Northern Turkey carried multiple copies of *cyp51* and the Y136F mutation. This suggests that (partial) resistance to DMIs is widespread across the continent, including in southern countries. The available data on fungicide usage fits with this pattern, as DMIs have been widely adopted compared to other fungicides (Eurostat 2007).

Across pathogens several dozens of *cyp51* mutations have been reported, with different combinations providing resistance to different DMIs (Chartrain and Brown 2023; Cools et al. 2013; Glaab et al. 2024; Jørgensen et al. 2019; Leroux et al. 2007; Mohd-Assaad et al. 2016; Pereira et al. 2020; Vestergård et al. 2023). In Bgt we found additional mutations at high frequency, for example S79T and K175N. These two mutations may provide resistance to some DMIs, potentially in combination with Y136F. Alternatively, they could be compensatory mutations, mitigating the fitness cost of carrying Y136F in absence of fungicide pressure (Arnold et al. 2024; Meyers 2020), or they could also be neutral mutations that hitchhiked to high frequencies. Similarly, “heteroallelism” could be a mechanism to enhance insensitivity (Arnold et al. 2024). However, it could also have a compensatory effect in the absence of fungicides, or be a contingency of the evolutionary process without functional consequences.

Our finding that resistance to DMIs is likely to have emerged at least twice independently, confirms what was already observed for QoI resistance in *Zymoseptoria tritici* (Estep et al. 2015; Torriani et al. 2009). Beyond the two main groups (cluster **i** and **ii** in Figure 4), we found additional smaller clusters, which could represent multiple additional independent emergences of DMI insensitivity. However, isoRelate can only infer IBD segments up to *∼*25 sexual generations ago and DMIs were introduced in the 1970s. Therefore, multiple isolates may share the same most recent common ancestor at the *cyp51* locus, but if that common ancestor was over 25 generations ago, these isolates would be split in different clusters. This might be the case for clusters **i, v**, and **vi**, which have identical mutational profiles (Figures 4 and 5). Similarly, cluster **ii** and **vii** share the same mutations and tandem repeat unit. Whether these examples represent truly independent origins cannot be determined with this data. Finally, in cluster **iv** we found multiple examples of tandem repeats of different length within the same isolate, suggesting a more complex evolutionary history that could be investigated with long-read data.

Nevertheless, the main result of this analysis was that two major clusters became less sensitive to DMIs through different mutational pathways: more copies, shorter tandem repeat unit and homoallelic Y136F mutation for cluster **i**; fewer copies, longer tandem repeat unit and heteroallelic Y136F and S509T mutations for cluster **ii**. After their emergence they spread rapidly and independently over the continent in the last 20 to 30 years. This is indicative of how quickly new advantageous variants can expand in Europe and neighboring regions. If the selective pressure is sufficiently strong, such as that imposed by fungicides and potentially also by resistance genes, new variants can increase in frequency and disperse over the continent within a few years.

### Other target genes

We found that there is little variation in the amino acid sequences and copy number for the three subunits of the succinate dehydrogenase gene (*sdhB, sdhC*, and *sdhD*), *erg2* and the beta tubulin genes. For beta tubulin this is expected, as MBC fungicides have fallen out of use in Europe (Jørgensen et al. 2019). Conversely, both morpholines and SDHIs are commonly applied on wheat fields, and at least in theory these genes should show signatures of selection. While *erg2* is the target of some morpholines (together with *erg24*), no resistance mutations were reported so far in Bgt. However, different morpholines can have different targets (*erg2* or *erg24*; Chartrain and Brown 2023), and it is possible that the main products applied on wheat fields in Europe do not target *erg2*. Similarly, the succinate dehydrogenase gene subunits are expected to be under selective pressure due to SDHI applications. Cereal powdery mildews might have an intrinsically low sensitivity to this fungicide class (Jørgensen et al. 2019), and this could explain the absence of resistance mutations in Bgt. However, mutations in *sdh* associated with resistance have been identified in grapevine powdery mildew (Stergiopoulos et al. 2022). Importantly, with our data we could not investigate the expression level of these genes, which could be linked to insensitivity to fungicides and be under selection.

## Conclusions

In this study we highlighted the potential of genomics for monitoring fungicide resistance. With our data we could reveal the landscape of fungicide resistance in European populations of Bgt. Beyond screening for resistance mutations and copy number variation, genomics allowed us to reconstruct the evolutionary history of different fungicide targets, something that would not be possible with sensitivity screenings. Continuous sampling and whole genome sequencing of these populations would be valuable, as they could reveal the early emergence of new mutations and their spread. This type of data would be exceptionally useful if coupled with sensitivity assays and gene expression analyses, as it could disentangle the effect of different mutations and variation in copy number. Moreover, long-read data would help in resolving complex loci such as *cyp51*. Finally, for a more informed interpretation of these results it would be important to have up-to-date and granular information about fungicide usage in different European regions, that could be correlated with the observed patterns of diversity in target genes and the epidemiological dynamics of resistance mutations.

## Experimental procedures

### WGS dataset

The data used in this study was generated by a previous study on the population genetics of wheat powdery mildew in Europe, which included 415 Bgt samples collected in Europe and surrounding regions between 1980 and 2023 (dataset *Europe+* in Jigisha et al. 2024). We based all our analyses on three sets of strains: 1) all isolates in the Europe+ dataset, 2) a subset of isolates collected in 2015 or more recently (dataset *Europe+_recent*), and 3) a subset of 83 samples collected over 40 years from the UK, France and Switzerland (*temporal* dataset). For all isolates included in the dataset accession numbers for the short reads are available in Supplementary Data S1.

### Fungicide targets

We focused on eight known fungicides targets which we also reported in Table 1. Three subunits of the succinate dehydrogenase gene (*sdhB, sdhC*, and *sdhD*), the sterol 14-demethylase gene (*cyp51*), the beta tubulin gene (*Btub*), cytochrome b (*cytb*), the C-8 sterol isomerase gene (*erg2*), and the delta(14)-sterol reductase (*erg24*).

For the 11 nuclear chromosomes we obtained the annotation from Müller et al. 2019 (PRJEB28180), while for the mitochondrial genome we used the annotation of Zaccaron and Stergiopolulos (2021; MT880591). We used homology with the corresponding genes of *Erysiphe necator* and RNAseq data to check the quality of the annotation. More specifically we used STAR v2.7.11b with default parameters (Dobin et al. 2013) to map RNAseq reads for two Bgt isolates (the reference strain 96224 and 94202; Praz et al. 2018; GEO accession number GSE108405), and we used IGV v2.16.2 (Thorvaldsdóttir et al. 2013) to visualize the read alignments. We updated the annotation of *sdhB* and *sdhC* (extending the N-terminus), we annotated *sdhD*, which was absent, and we left unchanged the other targets. The genomic coordinates for all gene targets are available at https://github.com/fmenardo/Bgt_fungicides_2024/tree/main/Fungicide_targets.

### Bioinformatic pipeline

We used the same pipeline used in Jigisha et al. 2024. Briefly, raw sequence reads were trimmed using fastp v0.23.2 (Chen 2023) with options with options cut_front_window_size 1, cut_front_mean_quality 20, cut_right_window_size 5, cut_right_mean_quality 20, –merge, overlap_len_require = 15, overlap_diff_percent_limit = 10, –cut_front, and –cut_right. Adapters were trimmed based on per-read overlap analysis employing the default settings in fastp. All resulting reads were mapped separately using bwa-mem (Li 2013) to the reference genome 96224 (Jigisha et al. 2024, Müller et al. 2019, Zaccaron & Stergiopoulos 2021). The alignments produced by bwa-mem were sorted and merged using samtools v1.17 (Danecek et al. 2021). Placeholder read-group and library information was added to the alignment files to make them compatible with GATK v.4.4.0 (Van der Auwera & O’Connor 2020). Duplicate reads were marked using GATK MarkDuplicatesSpark. Sample-level haplotype calling was performed with GATK HaplotypeCaller with options –ploidy 1 –ERC BP_RESOLUTION to produce a VCF file with calls for each site in the genome. The single VCF files of all samples were merged using GATK CombineGVCFs. GATK GenotypeGVCFs was used to perform joint genotyping on all the samples. Site-level hard filtering was executed using GATK VariantFiltration with filters QD<10, FS>55, MQ<45 and –4<ReadPosRankSum<4. The code for all steps of this analysis can be found at https://github.com/fmenardo/Bgt_popgen_Europe_2024/blob/Bgt_ms/WGS_pipeline/WGS_pipeline.md.

### Copy number variation

We calculated the average coverage for the nuclear genome (the 11 chromosomes) and for the mitochondrial genome with samtools coverage v1.17 (Danecek et al. 2021). We used the same approach to calculate the coverage for all gene targets (from start to end). We used the ratio between the coverage of the gene targets and the average coverage of the nuclear or mitochondrial genome (depending on the gene target) to estimate the copy number for each sample. After confirming normality (Shapiro-Wilk *p*-value < 0.001), but not normality of variance (F test ratio of variances = 2.512, *p*-value = 0.0096), we performed a Welch two sample t-test to test for different number of *cyp51* copies in the temporal groups “1980-2001” and “2022-2023”. To study the structure of tandem repeat at the *cyp51* locus we plotted the ratio between the coverage of each base of the locus and the average genome-wide coverage. Plots and data processing were performed using R version 4.3 (R Core Team 2023) and the *tidyverse* package (Wickham 2023).

### Gene alignments

For each target gene we used bcftools v1.17 (Danecek et al. 2021) to generate vcf files for the coding sequence. In a second step we converted these files into fasta files, reverse complemented the sequences for genes on the reverse strand (*sdhB, sdhC, cyp51, erg2, erg24*, and *cytb*) and translated all sequences into proteins. To do this we used the standard genetic code, except for *cytb* for which we used the mold, protozoan, and coelenterate mitochondrial code (NCBI codon table 4; https://www.ncbi.nlm.nih.gov/Taxonomy/Utils/wprintgc.cgi?mode=c). In these alignments, sites with minor allele supported by more than 10% of reads were coded as missing data.

We found many “heterozygous” mutations in *cyp51* (i.e.; mutations with minor allele supported by more than 10% of reads), which was due to the presence of multiple gene copies. Therefore, we produced an additional gene alignment in which instead of coding “heterozygous” sites as missing we reported the alternate allele. In this alignment a SNP should be interpreted as present in at least one of the copies of *cyp51*, but not necessarily in all. All gene alignments are available at https://github.com/fmenardo/Bgt_fungicides_2024/tree/main/Alignments

### Analysis of *cyp51* locus on Pacbio assembly of CHVD042201

The Pacbio assembly of one of the isolates in our dataset (CHVD042201) was produced recently (Kunz et al. 2024; PRJNA1131794). We confirmed the copy number prediction of *cyp51* made with short read coverage using blastn v2.9.0+ (Camacho et al. 2009) and redotable v1.2 (http://www.bioinformatics.babraham.ac.uk/projects/redotable/).

### IsoRelate and haplotype networks

We used isoRelate (Henden et al. 2018) on a region of 3Mb around the *cyp51* locus (chromosome 8 from bp 4,000,000 to bp 7,000,000) to identify “identical-by-descent” (IBD) segments between pairs of samples in the *Europe+_recent* dataset. We selected all SNPs with no missing data and a minor allele frequency greater than 5%. Additionally, we excluded SNPs that could not be mapped unambiguously on the genetic map, which was obtained from Müller et al. 2019. We considered only IBD segments that were larger than 2cM, larger than 50 Kb, and with a minimum number of SNPs equal or greater than 50. We generated clusters by connecting isolates that were IBD over the coding sequence of *cyp51*, and we plotted the resulting graph with the R package igraph (https://cran.r-project.org/web/packages/igraph/index.html). The haplotype network of the nucleotide sequence of *erg24* was inferred with the R package pegas, using the parsimony algorithm (haploNet function) (Paradis et al. 2010).

## Data availability

The accession numbers for short read sequence data used in this study are available in Supplementary Data S1. Code and additionla data to reproduce all analyses is available at https://github.com/fmenardo/Bgt_fungicides_2024. All other data is contained within the manuscript and its Supporting Information.

## Acknowledgements

This work was funded by the Swiss National Science Foundation grant PZ00P3_193473. We would like to thank Daniel Croll, Matthias Heuberger, and Thomas Wicker for their feedback on tandem repeats. Luca Cornetti and Stefano F.F. Torriani work or have worked for the fungicide industry (Syngenta Crop Protection AG). The rest of the authors declare no conflict of interest.

