## Supplementary Information for "Genomic surveillance and molecular evolution of fungicide resistance in European populations of wheat powdery mildew"

Minadakis et al.

### Tables

**Table S1. All amino acid mutations in the eight target genes**
The odds ratio and the $p$-values refer to the Fisher's exact test on the temporal dataset. For mutations not occurring in France, Switzerland and the UK the test could not be performed.

| Gene target | Mutation | N. of isolates | Odds ratio | $p$-value |
|---|---|---|---|---|
| *cytb* | G143A | 146 | 0.04769099 | 4.378 e-6 |
| *Btub* | - | - | - | - |
| *erg2* | F29I | 1 | - | - |
| *erg2* | V59I | 1 | - | - |
| *erg2* | K89N | 5 | - | - |
| *erg2* | L115I | 1 | - | - |
| *erg2* | N177D | 3 | - | - |
| *sdhB* | A14T | 1 | - | - |
| *sdhB* | E36K | 3 | - | - |
| *sdhB* | A263S | 1 | - | - |
| *sdhC* | S9F | 9 | - | - |
| *sdhC* | S10L | 1 | - | - |
| *sdhC* | P29L | 1 | - | - |
| *sdhC* | F35I | 2 | - | - |
| *sdhC* | S38Y | 3 | - | - |
| *sdhC* | A130S | 1 | - | - |
| *sdhD* | T5I | 1 | - | - |
| *sdhD* | N22K | 1 | - | - |
| *sdhD* | H34N | 1 | - | - |
| *sdhD* | A90V | 1 | - | - |
| *sdhD* | A137V | 1 | - | - |
| *erg24* | D137E | 17 | - | - |
| *erg24* | Y165F | 74 | 1.337203 | 0.732 |
| *erg24* | H239N | 1 | - | - |
| *erg24* | F289H | 36 | 0 | 0.1947 |
| *erg24* | D291N | 8 | - | - |
| *erg24* | V295L | 141 | 0.3555229 | 0.07493 |
| *erg24* | F316F | 1 | - | - |
| *erg24* | L357F | 1 | - | - |
| *cyp51* | I3K | 19 | - | - |
| *cyp51* | S79T | 211 | 0.1619583 | 0.002381 |
| *cyp51* | Y136F | 298 | 0.02111874 | 1.024e-05 |
| *cyp51* | K175N | 234 | 0.1133233 | 0.0006171 |
| *cyp51* | L236F | 14 | - | - |
| *cyp51* | T271S | 14 | - | - |
| *cyp51* | E438K | 1 | - | - |
| *cyp51* | S509T | 62 | 0 | 0.03079 |

### Figures

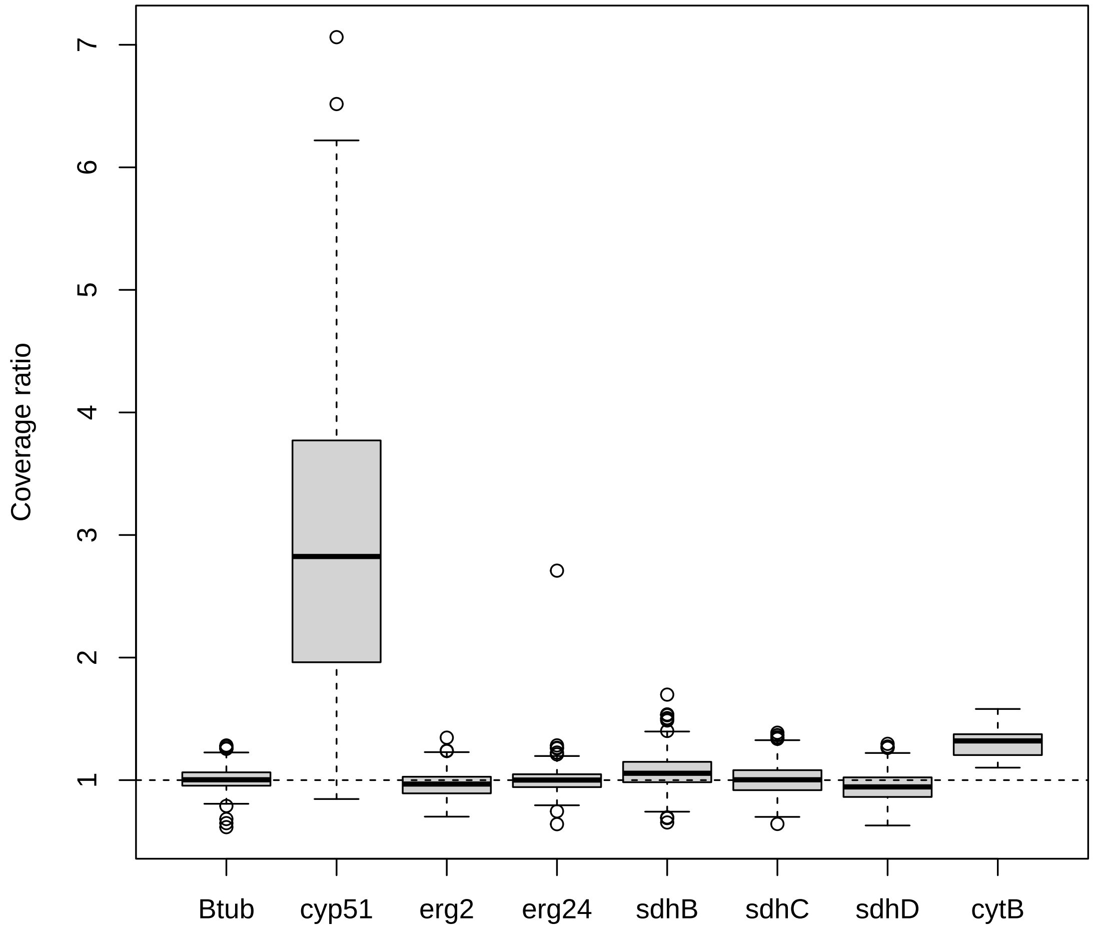

**Figure S1. Coverage ratio for the eight studied genes**
The ratios of gene coverage and genome-wide coverage were used to estimate the number of copies for each gene. For *cytb* we used the mitochondrial genome coverage as denominator.

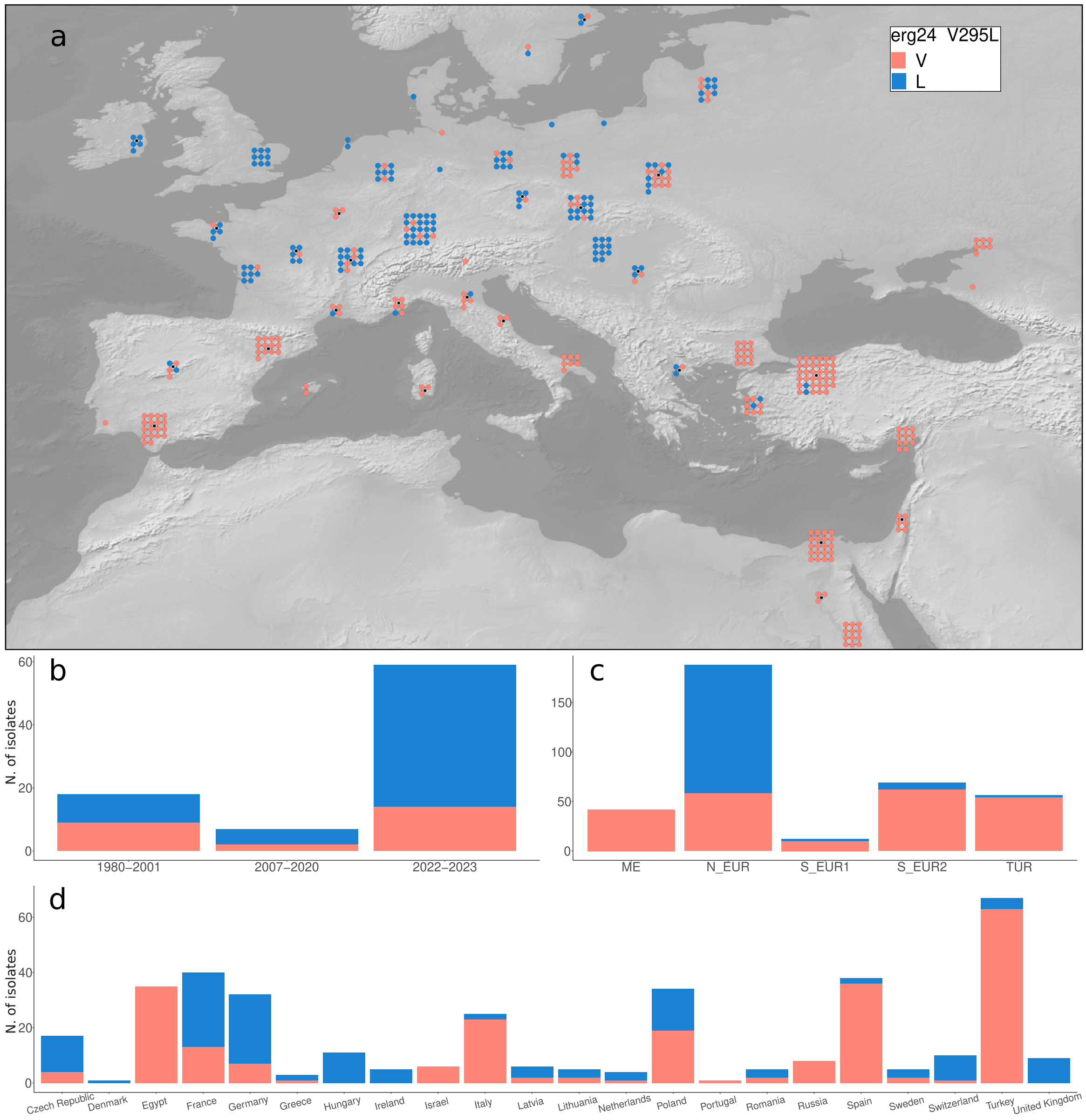

**Figure S2.** ***erg24*** **mutations V295L**
**(a)** Distribution of V295L. **(b)** Frequency of V295L by year of collection (*temporal* dataset). **(c)** Frequency of V295L by population. **(d)** Frequency of V295L by country of origin.

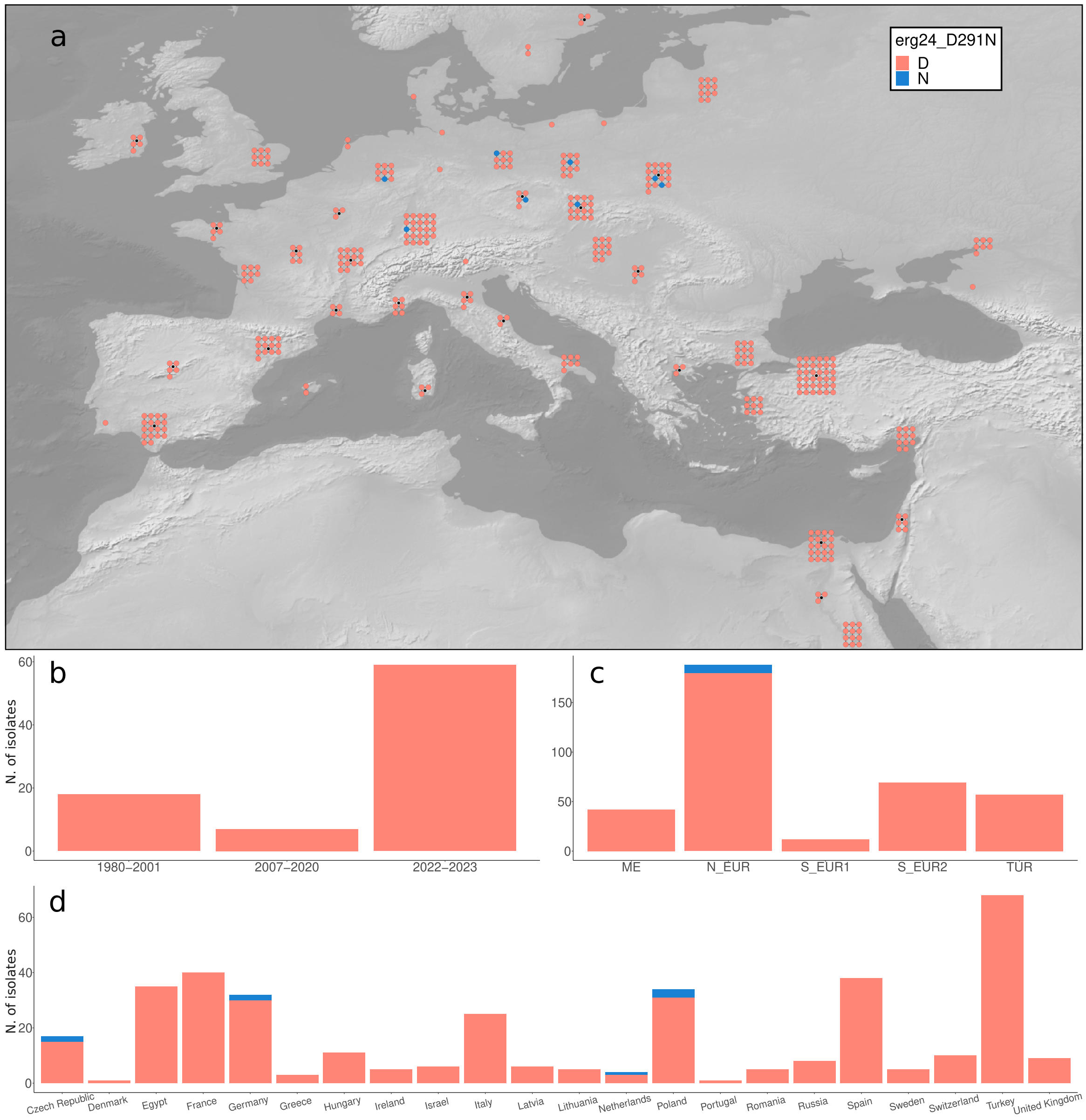

**Figure S3.** ***erg24*** **mutation D291N**
**(a)** Distribution of D291N. **(b)** Frequency of D291N by year of collection (*temporal* dataset). **(c)** Frequency of D291N by population. **(d)** Frequency of D291N by country of origin.

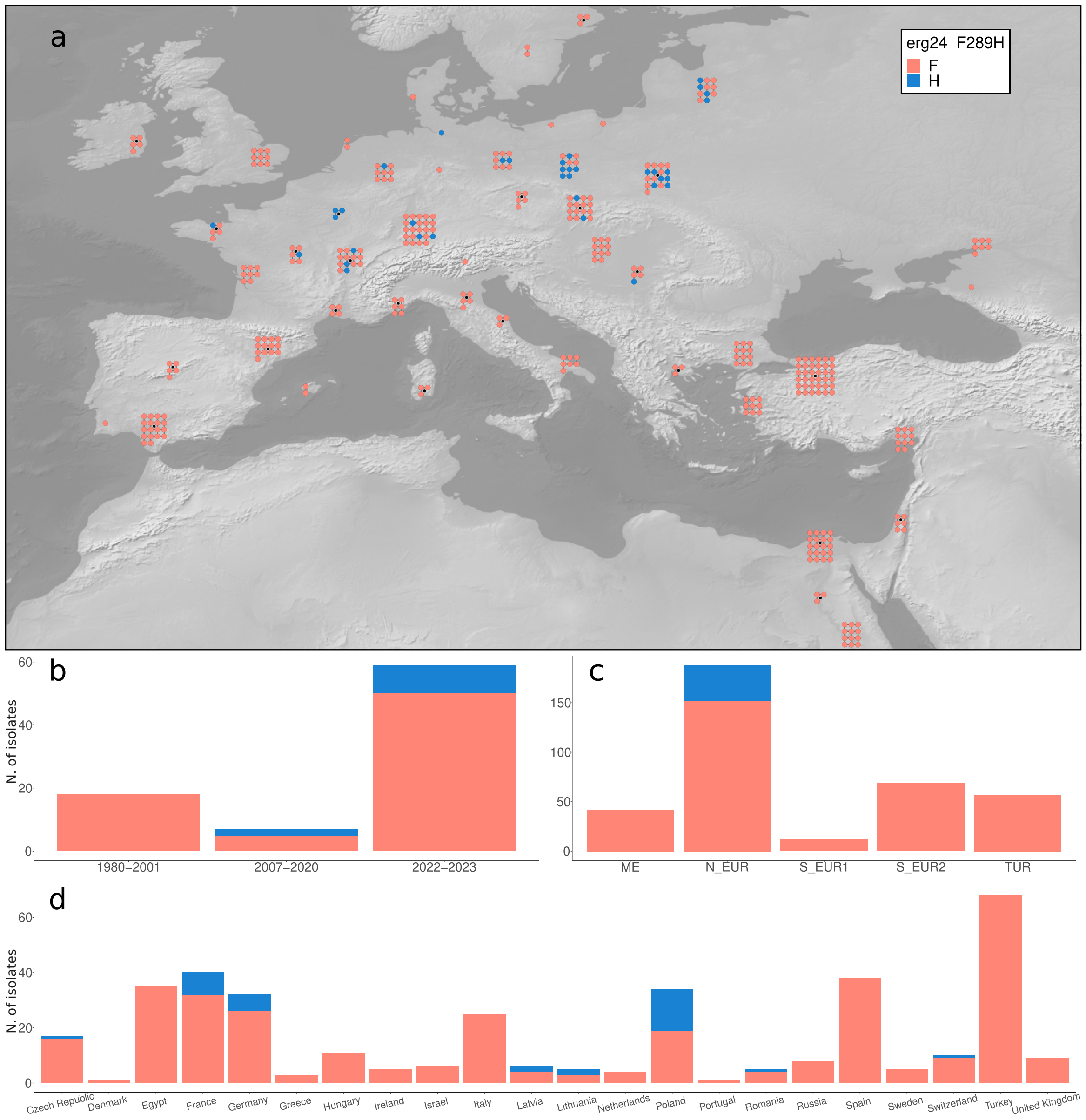

**Figure S4.** ***erg24*** **mutation F289H**
**(a)** Distribution of F289H. **(b)** Frequency of F289H by year of collection (*temporal* dataset). **(c)** Frequency of F289H by population. **(d)** Frequency of F289H by country of origin.

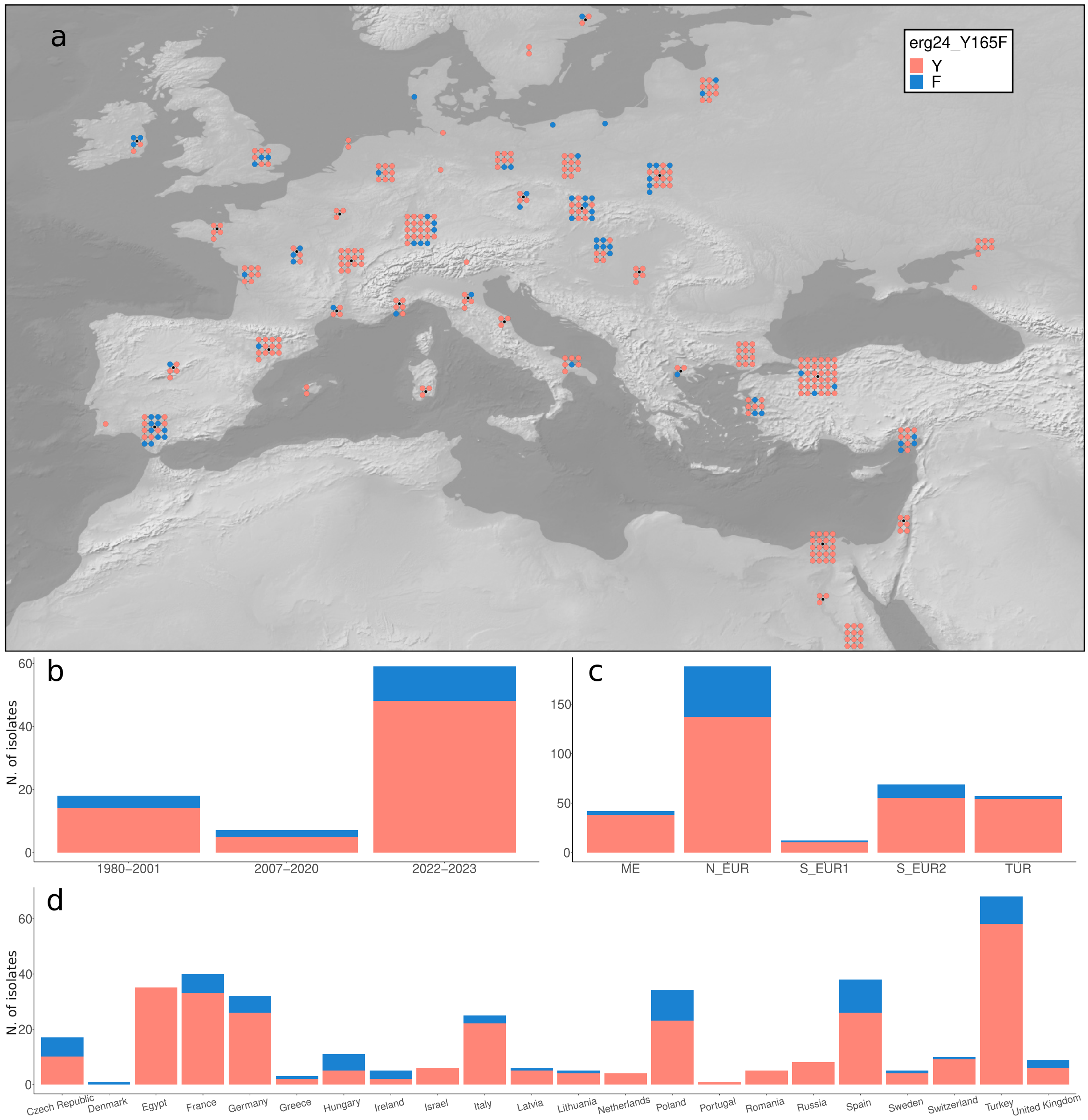

**Figure S5.** ***erg24*** **mutation Y165F**
**(a)** Distribution of Y165F. **(b)** Frequency of Y165F by year of collection (*temporal* dataset). **(c)** Frequency of Y165F by population. **(d)** Frequency of Y165F by country of origin.

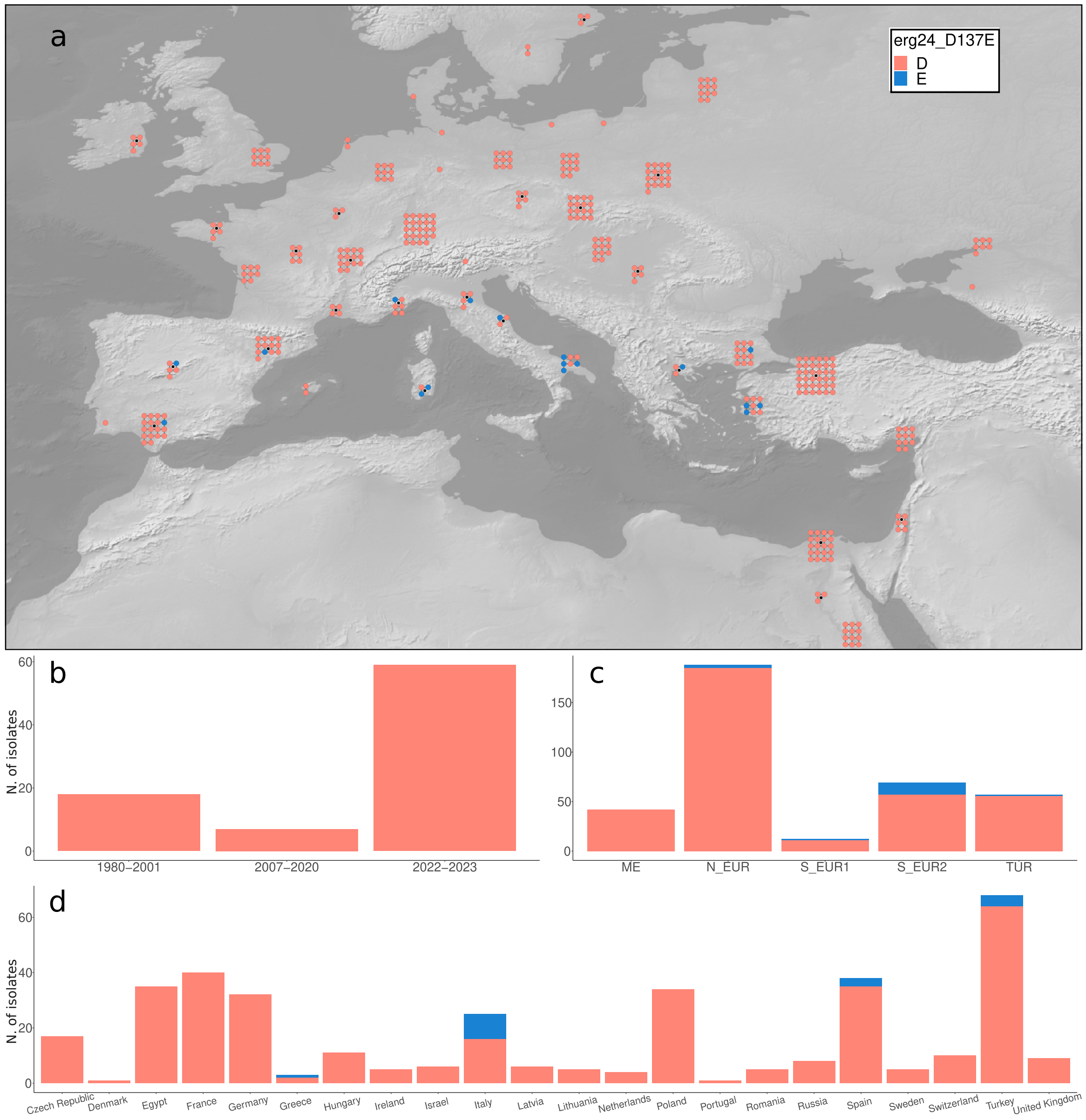

**Figure S6.** ***erg24*** **mutation D137E**

**(a)** Distribution of D137E. **(b)** Frequency of D137E by year of collection (*temporal* dataset). **(c)** Frequency of D137E by population. **(d)** Frequency of D137E by country of origin.

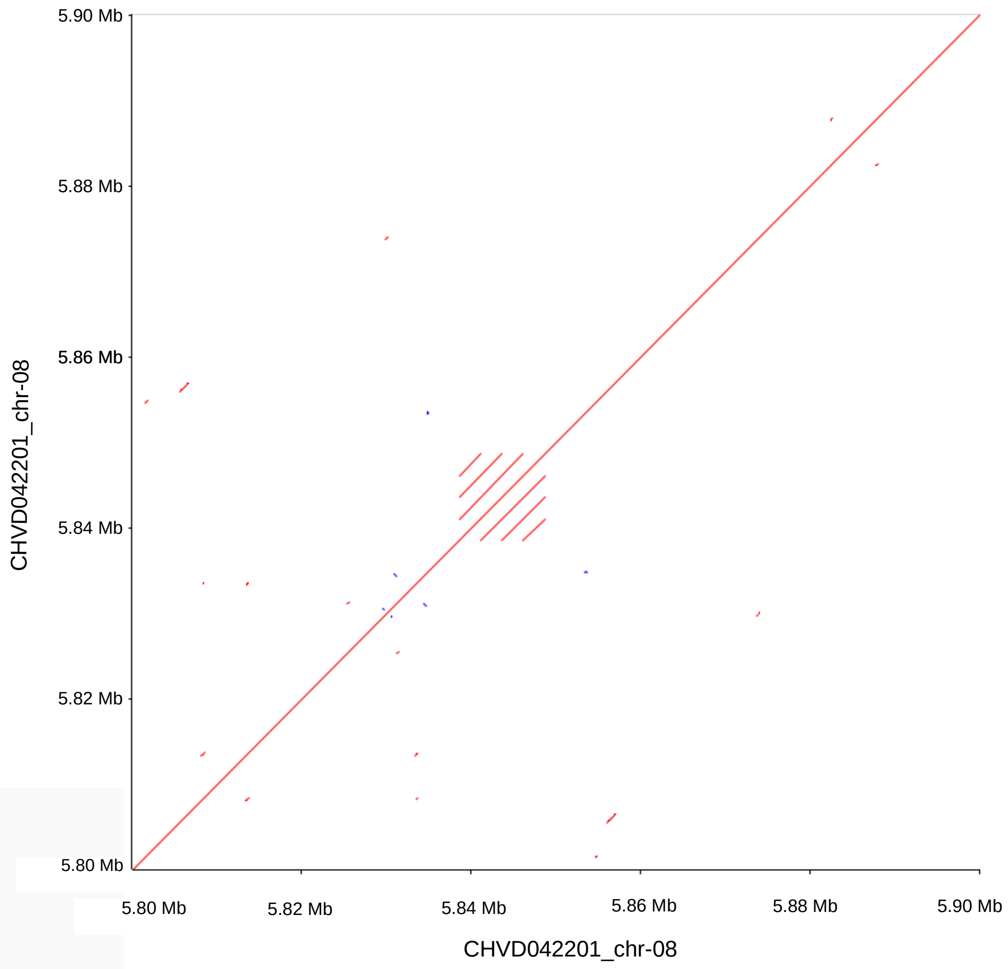

**Figure S7. Dotplot of *cyp51* locus for isolate CHVD042201** Dotplot of 100Kb of chromosome 8 in correspondence of the *cyp51* locus. The same genomic fragment is plotted on the x and y axes. Four copies of *cyp51* are located in the tandem repeat.

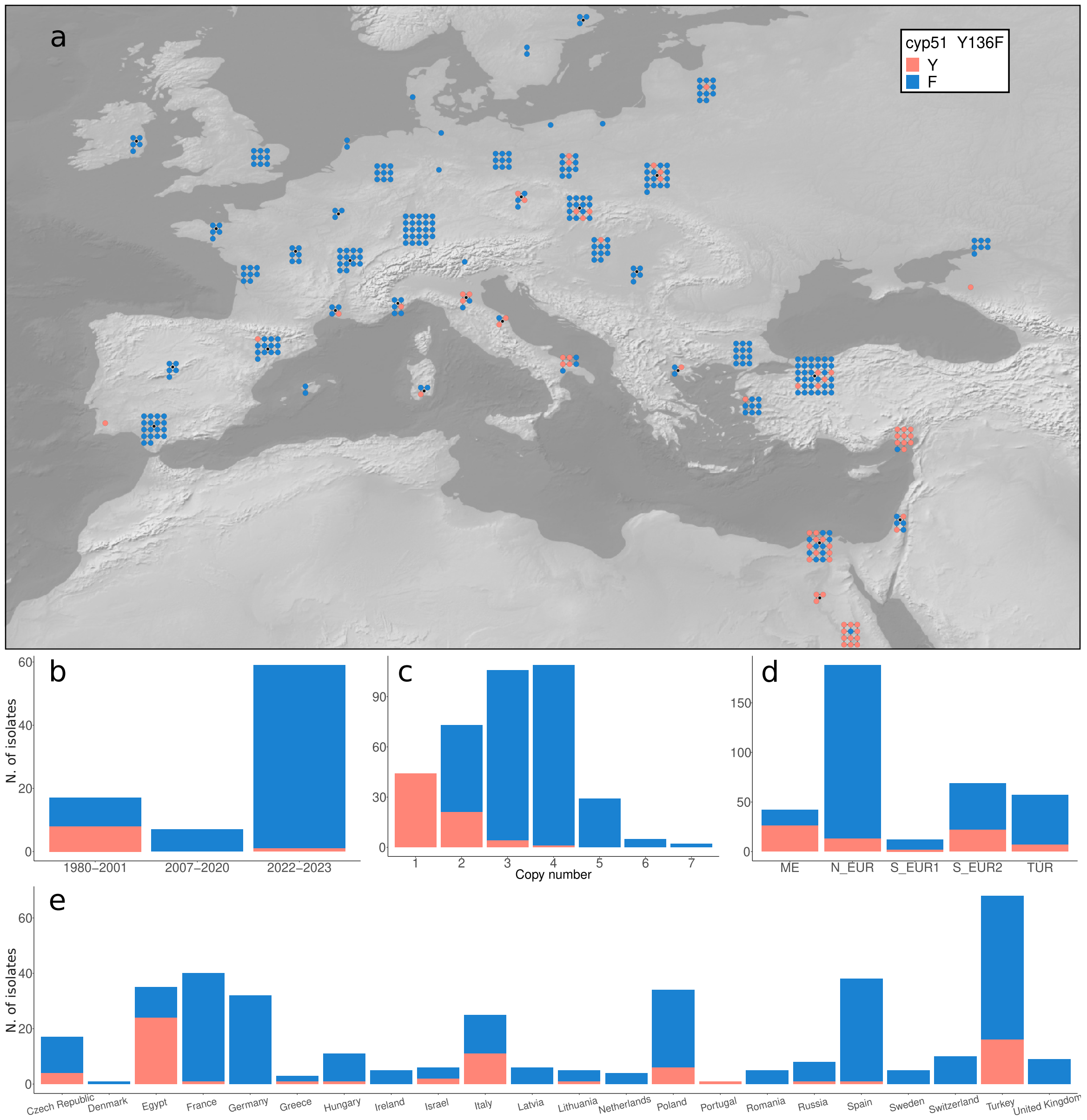

**Figure S8.** ***cyp51*** **mutation Y136F**
**(a)** Distribution of Y136F. **(b)** Frequency of Y136F by year of collection (*temporal* dataset). **(c)** Frequency of Y136F by population. **(d)** Frequency of Y136F by country of origin.

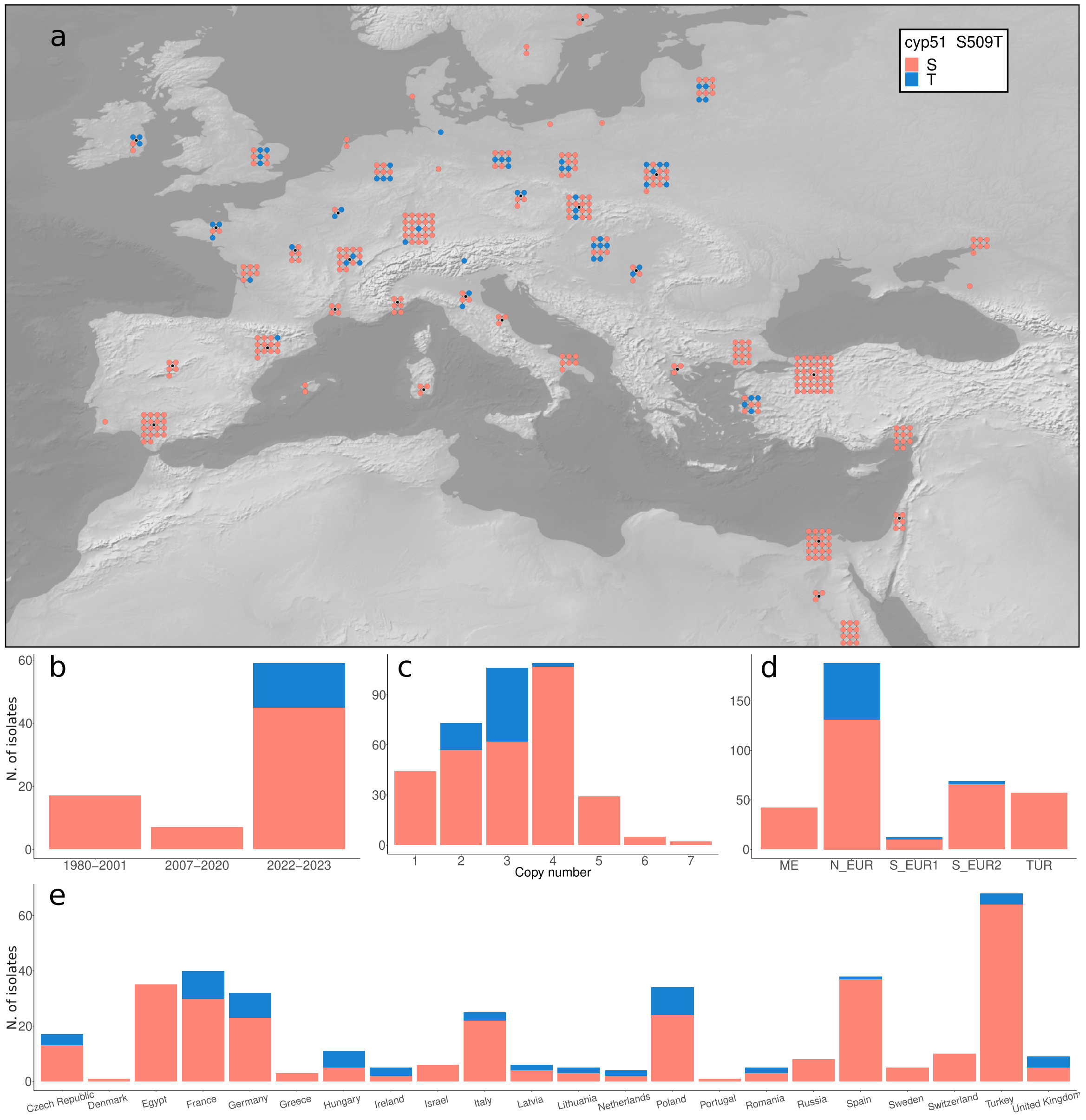

**Figure S9.** ***cyp51*** **mutation S509T**
**(a)** Distribution of S509T. **(b)** Frequency of S509T by year of collection (*temporal* dataset). **(c)** Frequency of S509T by population. **(d)** Frequency of S509T by country of origin.

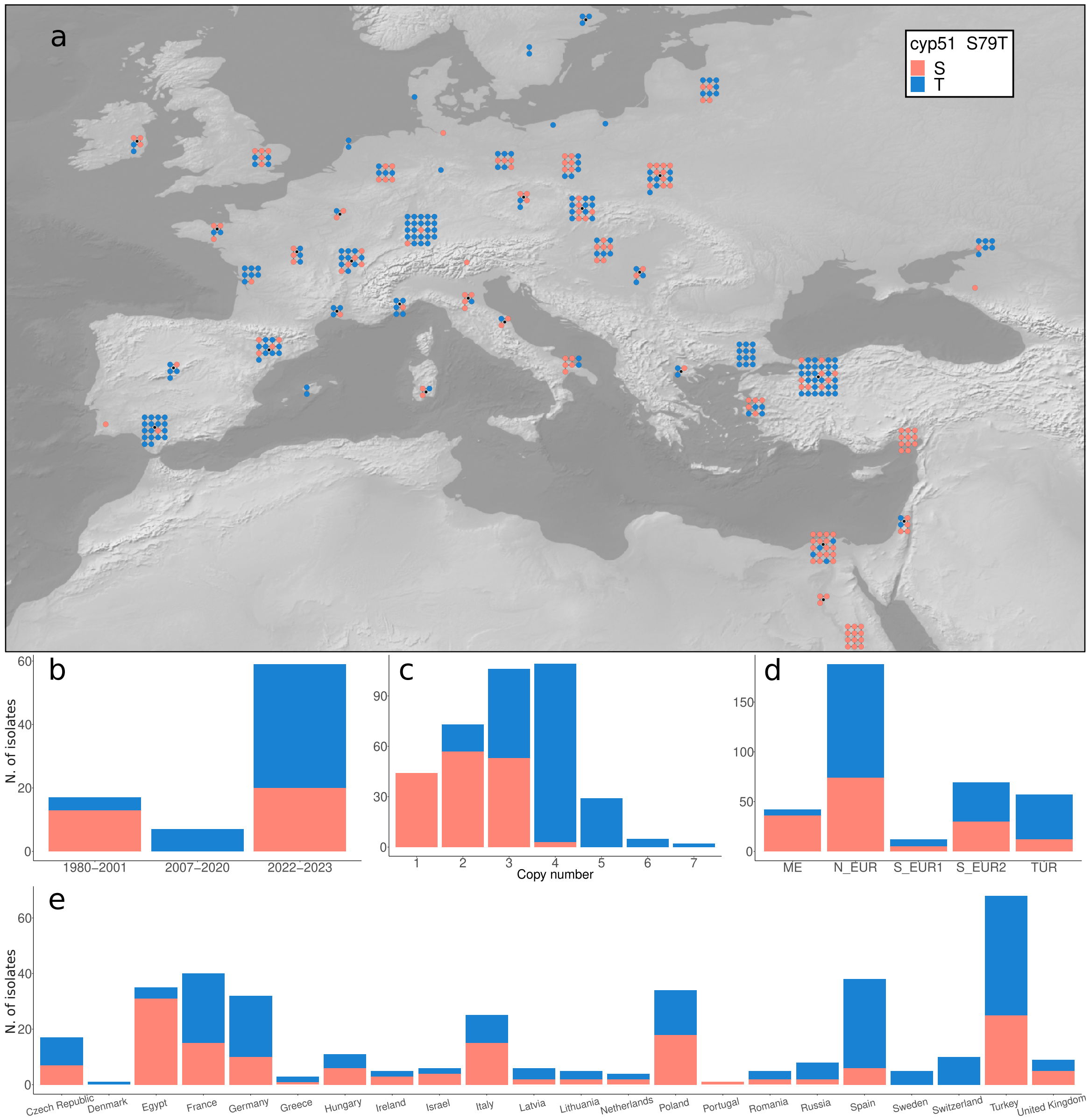

**Figure S10.** ***cyp51*** **mutation S79T**

**(a)** Distribution of S79T. **(b)** Frequency of S79T by year of collection (*temporal* dataset). **(c)** Frequency of S79T by population. **(d)** Frequency of S79T by country of origin.

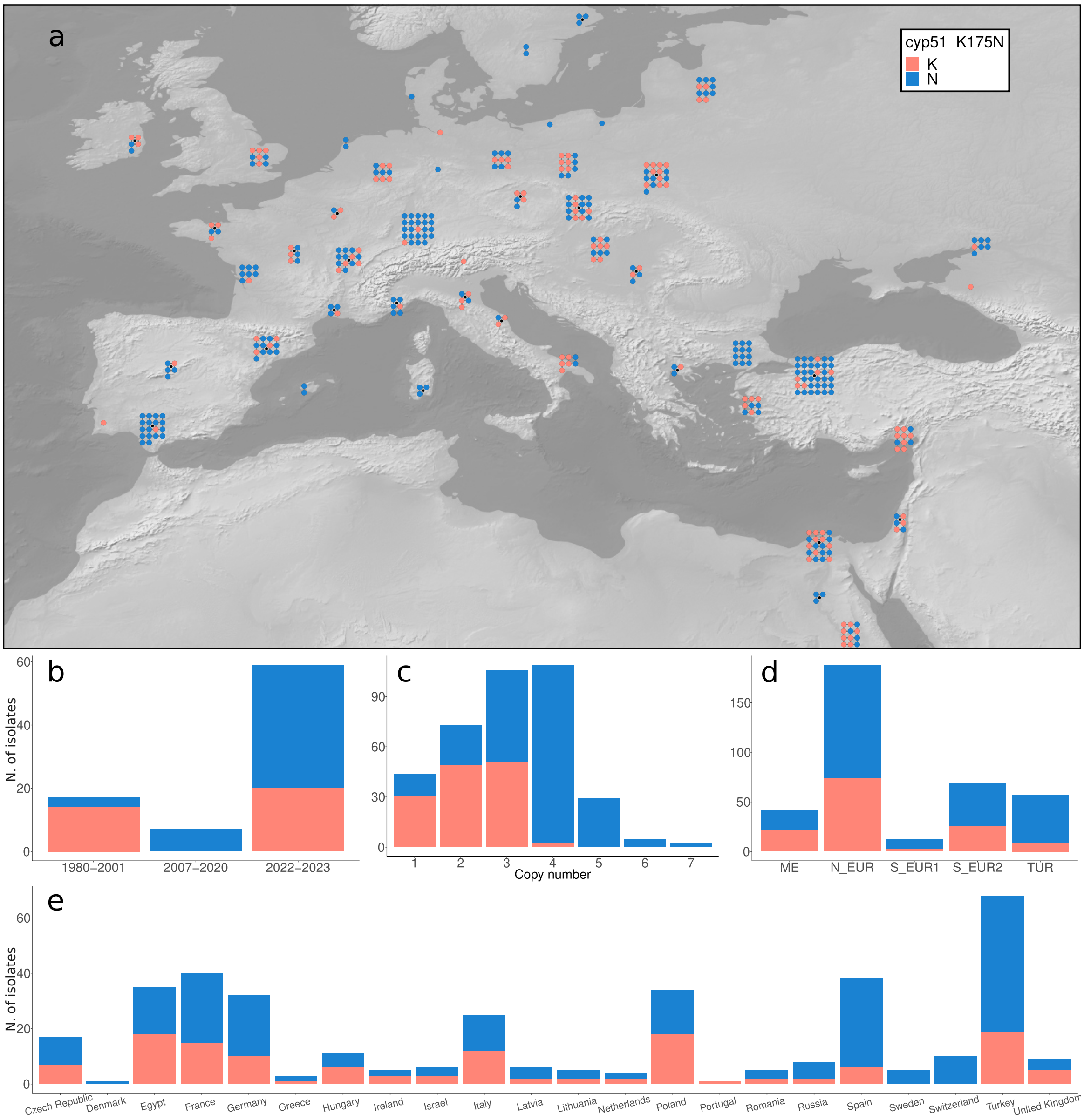

**Figure S11.** ***cyp51*** **mutation K175N**
**(a)** Distribution of K175N. **(b)** Frequency of K175N by year of collection (*temporal* dataset). **(c)** Frequency of K175N by population. **(d)** Frequency of K175N by country of origin.

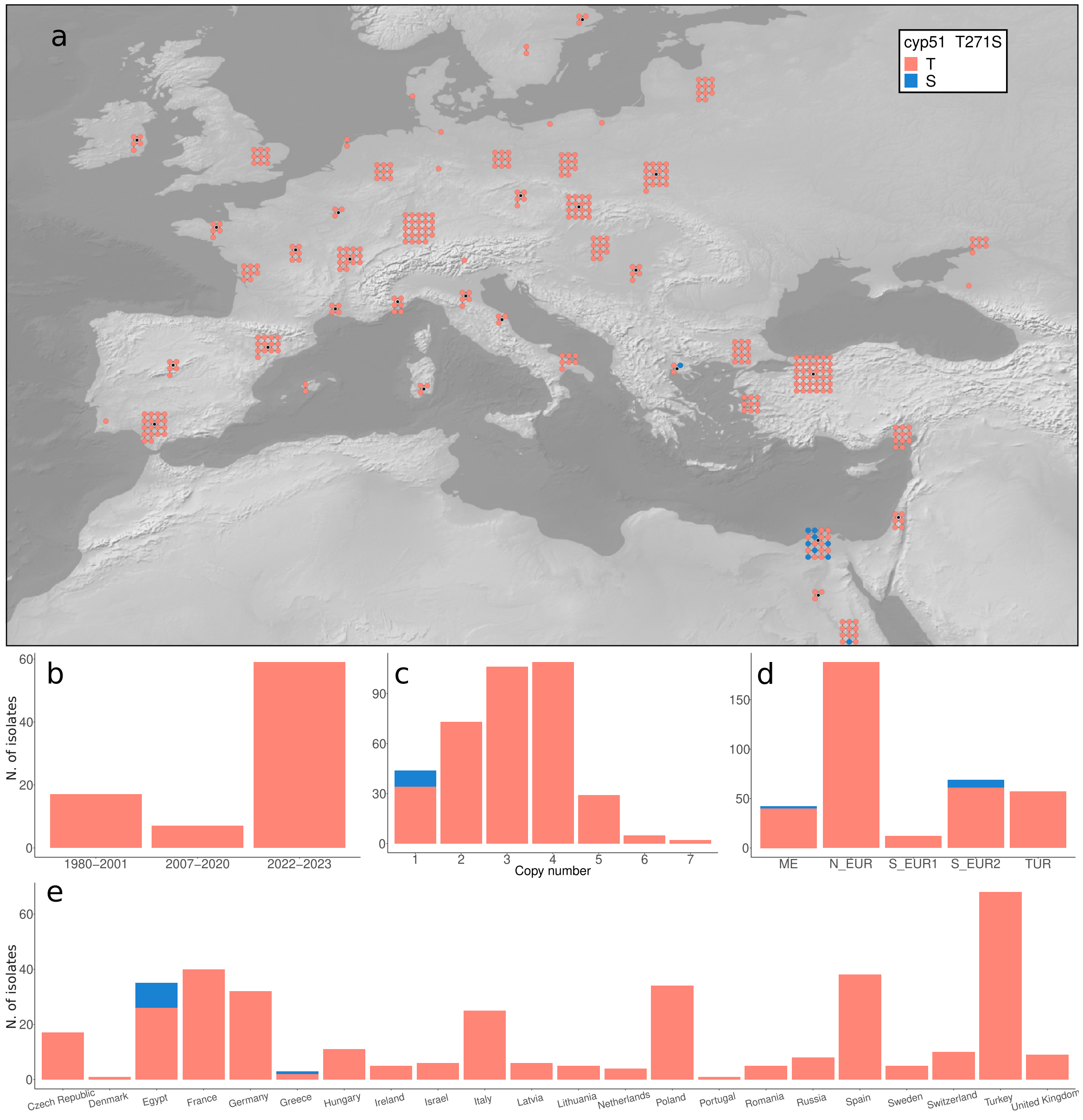

**Figure S12.** ***cyp51* mutation T271S**

**(a)** Distribution of T271S. **(b)** Frequency of T271S by year of collection (*temporal* dataset). **(c)** Frequency of T271S by population. **(d)** Frequency of T271S by country of origin.

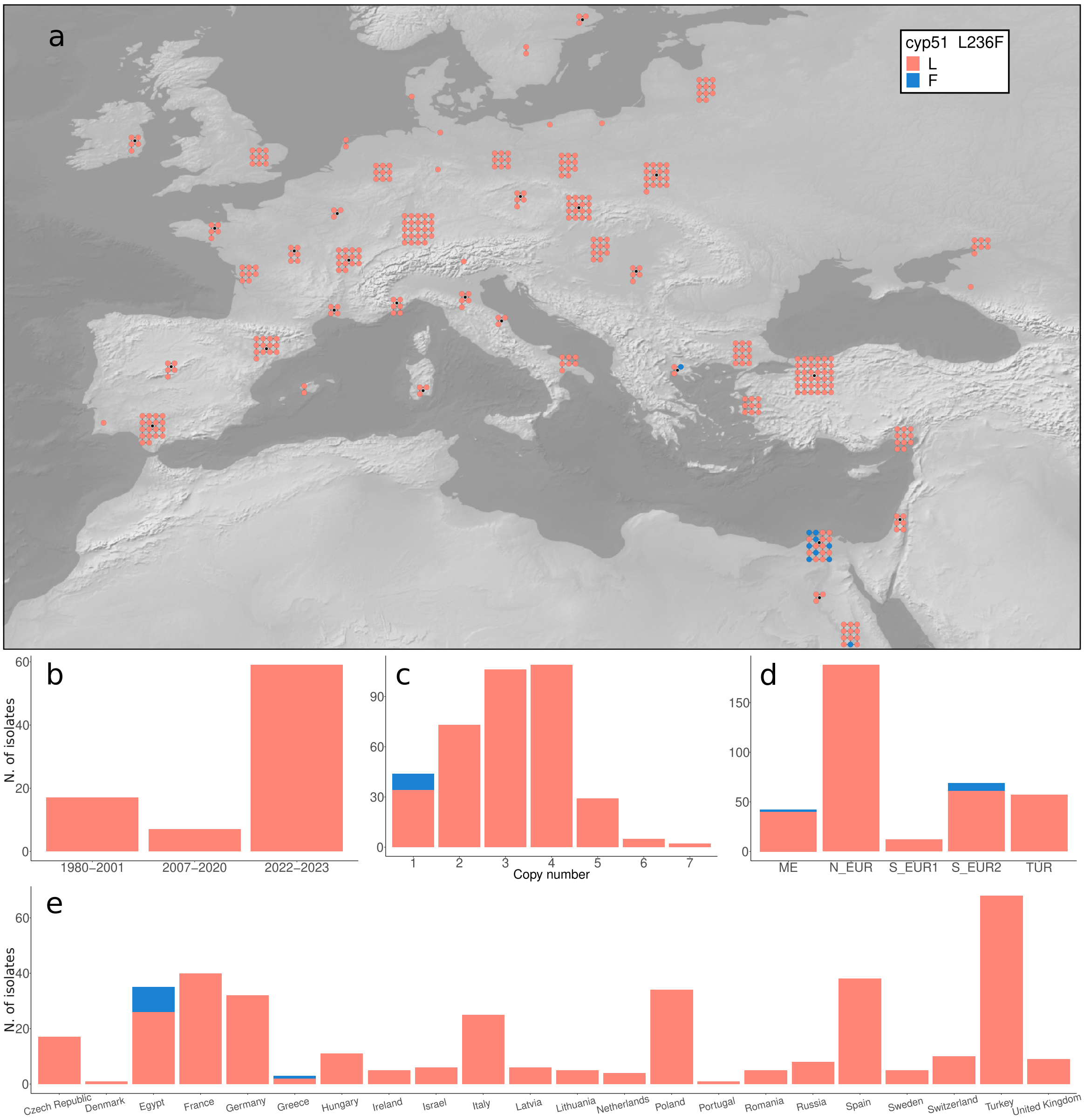

**Figure S13.** ***cyp51*** **mutation L236F**
**(a)** Distribution of L236F. **(b)** Frequency of L236F by year of collection (*temporal* dataset). **(c)** Frequency of L236F by population. **(d)** Frequency of L236F by country of origin.

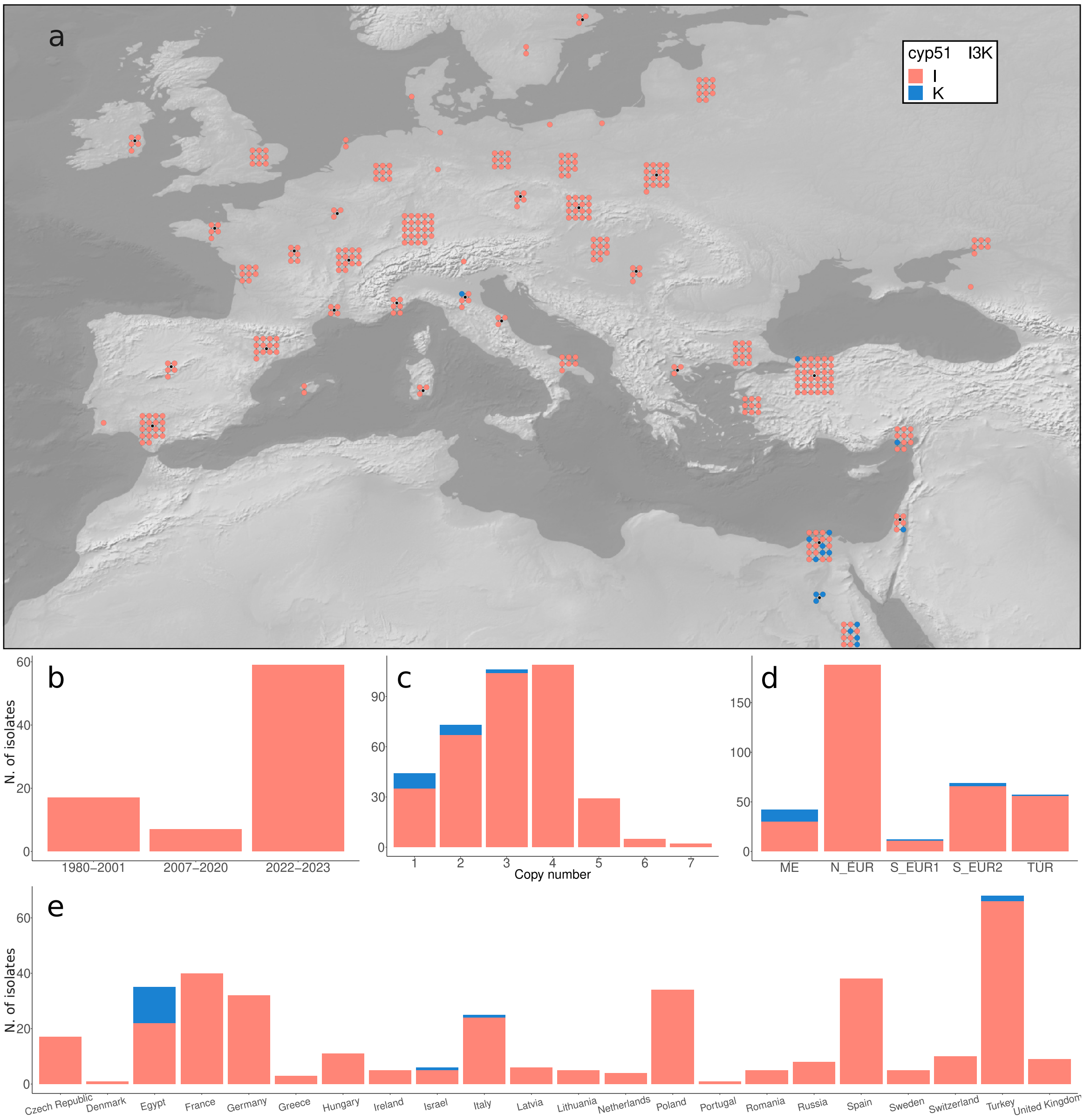

**Figure S14. *cyp51* mutation I3K**
**(a)** Distribution of I3K. **(b)** Frequency of I3K by year of collection (*temporal* dataset). **(c)** Frequency of I3K by population. **(d)** Frequency of I3K by country of origin.
